# A TRPV4-dependent calcium signaling axis governs lamellipodial actin architecture to promote cell migration

**DOI:** 10.1101/2025.03.28.646012

**Authors:** Ernest Iu, Alexander Bogatch, Wenjun Deng, Jonathan D. Humphries, Changsong Yang, Fernando R. Valencia, Chengyin Li, Christopher A. McCulloch, Guy Tanentzapf, Tatyana M. Svitkina, Martin J. Humphries, Sergey V. Plotnikov

## Abstract

Cell migration is crucial for development and tissue homeostasis, while its dysregulation leads to severe pathologies. Cell migration is driven by the extension of actin-based lamellipodia protrusions, powered by actin polymerization, which is tightly regulated by signaling pathways, including Rho GTPases and Ca^2+^ signaling. While the importance of Ca^2+^ signaling in lamellipodia protrusions has been established, the molecular mechanisms linking Ca^2+^ to lamellipodia assembly are unknown. Here, we identify a novel Ca^2+^ signaling axis involving the mechano-gated channel TRPV4, which regulates lamellipodia protrusions in various cell types. Using Ca^2+^ and FRET imaging, we demonstrate that TRPV4-mediated Ca^2+^ influx upregulates RhoA activity within lamellipodia, which then facilitates formin-mediated actin assembly. Mechanistically, we identify CaMKII and TEM4 as key mediators relaying the TRPV4-mediated Ca^2+^ signal to RhoA. These data define a molecular pathway by which Ca^2+^ influx regulates small GTPase activity within a specific cellular domain – lamellipodia - and demonstrate the critical role in organizing the actin machinery and promoting cell migration in diverse biological contexts.

## INTRODUCTION

Cell migration underlies a diverse array of physiological processes, ranging from embryonic development to tissue homeostasis. When cell migration is dysregulated, devastating pathological conditions often arise, including developmental defects, chronic wounds and immunodeficiencies^1^.

Over the past decades, significant progress has been made in understanding the molecular mechanisms that govern cell migration. It is now well established that cells rely on various subcellular compartments, protein assemblies, and organelles to facilitate and coordinate movement^2^. At the leading edge of a migrating cell, dynamic membrane protrusions extend in the direction of movement to propel the cell forward. In mesenchymal cells, such as fibroblasts and metastatic cancer cells, lamellipodia are the primary machinery that enables rapid migration through complex biological microenvironments^3,4^. Lamellipodia are formed by the polymerization of globular actin (G-actin) into filamentous actin (F-actin) with the aid of the Arp2/3 complex and formins^5,6^. The polymerizing actin filaments elongate and push against the plasma membrane, providing the biophysical basis of membrane protrusion^7^. For efficient cell movement, the dynamics of actin polymerization must be spatially and temporally regulated^8^. Disruption of signaling pathways essential for lamellipodial protrusions significantly impedes cell migration *in vitro* and *in vivo*^9,10^.

The regulation of actin polymerization in lamellipodia has been largely attributed to Rho-family small GTPases, including RhoA, Rac1, and Cdc42, which play a central role in coordinating the spatiotemporal dynamics of the actin cytoskeleton in migrating cells^11^. These small GTPases are considered a major convergence point of many migration-related signaling pathways, such as phosphoinositides and receptor tyrosine kinases, for their ability to transduce multiple upstream signals to nucleation promoting factors, including the WAVE, and Arp2/3 complexes^12^. Among Rho-family GTPases, Rac1 is considered the main driver of lamellipodia formation due to its role in promoting branched actin formation by activating the Arp2/3 complex^13^. However, RhoA and Cdc42 are also activated in cellular protrusions and regulate actin assembly in lamellipodia^11,14–16^. It was proposed that RhoA initiates protrusions by activating the formin family of actin elongation factors, mDia1 and mDia2, while Rac1/Cdc42 activate other actin regulators, *e.g.*, the Arp2/3 complex and VASP, for continuous growth of the lamellipodia^5,17^.

In addition to small GTPases, calcium (Ca^2+^) ions have also emerged as potent regulators of cell migration. Previous studies have shown that Ca^2+^ signaling is implicated in various aspects of cell migration, from protrusion and adhesion formation to cell detachment^18–20^. A major effector of Ca^2+^ signaling is myosin-II, a molecular motor that governs the retraction of lamellipodia and facilitates the attachment of cell protrusions to the extracellular matrix (ECM) through focal adhesions^21–23^. In addition, several studies have demonstrated the importance of Ca^2+^ signaling in the regulation of actin dynamics at the cell edge and protrusive activity of lamellipodia^24,25^, but molecular players controlling Ca^2+^ signaling at the leading edge and transducing these signals to the dynamics of the lamellipodia actin network remain unknown.

Here, we elucidated a signaling axis that controls the organization of the lamellipodial actin network and protrusive activity of migrating cells. To determine the molecular basis of lamellipodia regulation by Ca^2+^ signaling, we sought to identify the primary source of Ca^2+^ that contributes to lamellipodia protrusion. By screening a panel of Ca^2+^-modulating drugs and siRNAs targeting Ca^2+^ channels in a cell spreading assay, we demonstrated that the channel activity of a mechano-gated channel TRPV4 is required for lamellipodia protrusions across different cell types. Ratiometric Ca^2+^ imaging demonstrated that suppression of TRPV4 disrupts Ca^2+^ influx in lamellipodia without perturbing the Ca^2+^ homeostasis in the cell body. We further demonstrated that suppression of TRPV4 decreases the activity of RhoA in lamellipodia and thereby reduces the density of lamellipodial F-actin. By using a combination of optogenetic perturbation, FRET imaging, and phospho-proteomic analysis, we identified two key molecular players — CaMKII (Ca^2+^/calmodulin-dependent protein kinase II) and its phosphorylation target, Tumour Endothelial Marker 4 (TEM4) — that transduce the Ca^2+^ signal from TRPV4 to RhoA. This Ca^2+^ signaling axis governs lamellipodia protrusions and cell migration in different biological contexts. Together, our findings reveal a conserved, local Ca^2+^ signaling axis composed of TRPV4, CaMKII, TEM4, and RhoA, that finetunes the actin machinery at the cell leading edge during cell migration.

## RESULTS

### Ca^2+^ influx through a mechano-gated channel TRPV4 facilitates lamellipodia protrusion

To investigate the role of Ca^2+^ signaling in the regulation of lamellipodia protrusion, we examined the intracellular Ca^2+^ dynamics of mouse embryonic fibroblasts (MEFs) undergoing cell spreading, a process aided by Arp2/3-containing lamellipodial actin networks (Figure S1A)^4,26^. MEFs were transfected with a construct encoding the Ca^2+^ biosensor GCaMP6f and soluble mCherry, separated by a T2A self-cleaving peptide, allowing near-equimolar expression of both proteins from a single mRNA transcript^27^.

Twenty-four hours post transfection, cells were allowed to spread on fibronectin (FN)- coated coverslips, and their spreading kinetics and Ca^2+^ patterns were visualized by time-lapse confocal microscopy. To determine the relative concentration of intracellular Ca^2+^, a ratiometric GCaMP6f signal was calculated by dividing the GCaMP6f intensity by the intensity of mCherry (hereafter referred to as the GCaMP signal). These experiments revealed a pronounced gradient of GCaMP signal across the cells that persisted throughout the entire duration of spreading and peaked at the protruding cell edge (Figure 1A). Furthermore, the local GCaMP signal intensity positively correlated with cell edge speed (Figure 1B), suggesting a functional link between Ca^2+^ signaling at the lamellipodia and its protrusive activity. Consistent with these findings, depletion of intracellular Ca^2+^ using BAPTA-AM abolished the GCaMP signal gradient and reduced cell spreading speed (Figures 1C-E and S1B). Conversely, cells treated with the Ca^2+^ ionophore ionomycin exhibited a rapid increase in GCaMP signal at the protruding cell edge, which coincided with a faster cell spreading (Figures S1C-E). Together, these data demonstrate an association between the local level of Ca^2+^ at the lamellipodia and cell protrusion.

**Figure 1.**
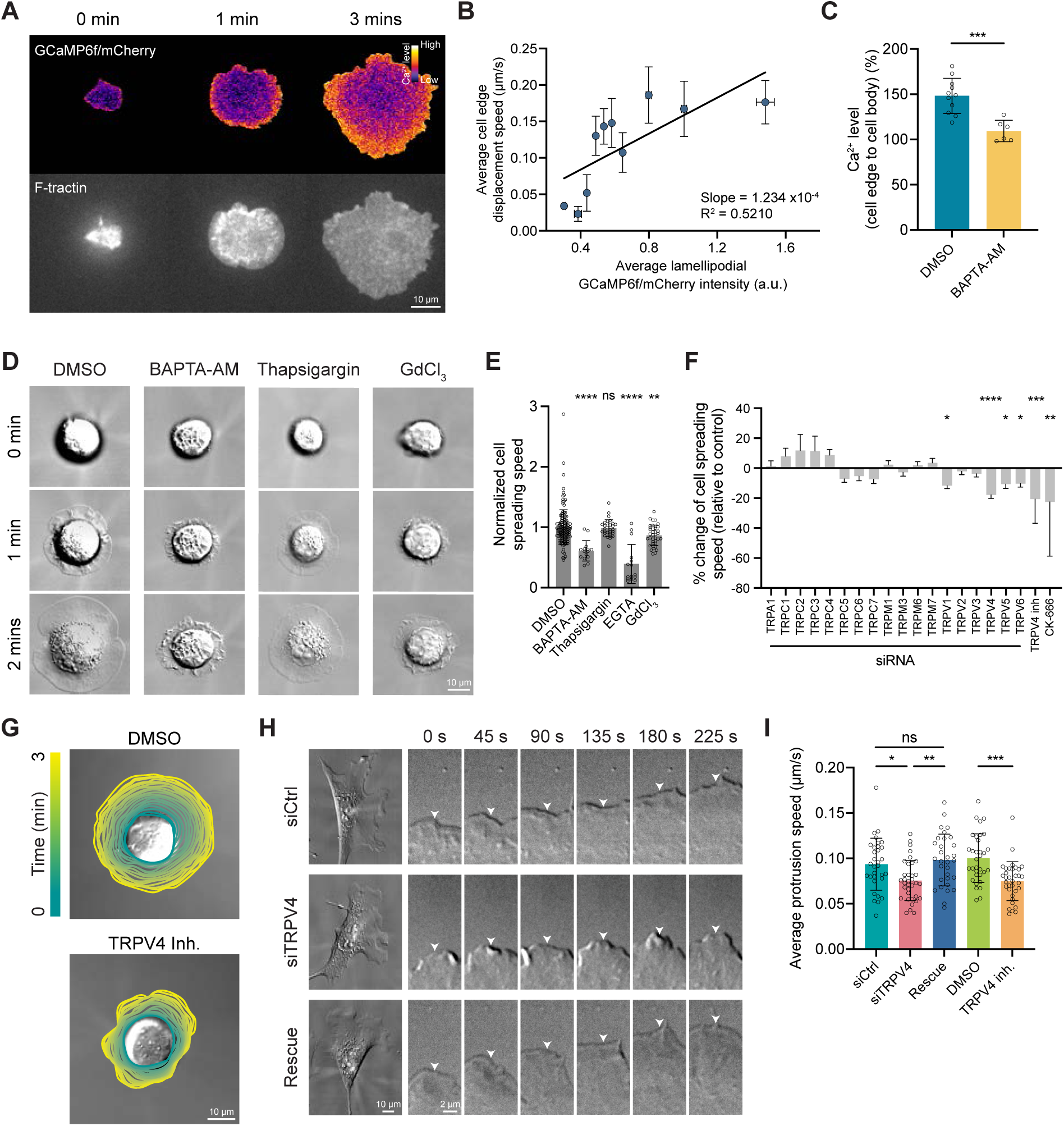
A mechano-gated Ca^2+^-permeable ion channel, TRPV4, regulates lamellipodial protrusions during cell spreading and migration. (A) Ratiometric GCaMP6f/mCherry and F-tractin-emiRFP670 images of a spreading MEF. (B) Quantification of lamellipodial GCaMP6f/mCherry intensity and cell edge displacement speed. Each dot represents data from a spreading cell. n = 10 cells. The error bars indicate the standard error (SE). (C) Quantification of the relative Ca^2+^ levels in cell edge compared to cell body in 0.1% DMSO- and 10 μM BAPTA-AM-treated cells in Ca^2+^-free DMEM. n ≥ 6 cells per condition. (D-E) Effects of Ca^2+^-perturbing drugs on cell spreading (D) DIC images of spreading cells treated with 0.1% DMSO, 10 μM BAPTA-AM, 10 μM GdCl3, or 1 μM thapsigargin. (E) Quantification of cell spreading speeds. Each data point is normalized to the mean of its control condition. n ≥ 15 cells per condition. (F) Quantification of cell spreading speeds upon siRNA depletion of individual TRP family channels, TRPV4 inhibition (10 μM HC-067047), or Arp2/3 inhibition (100 μM CK-666). n ≥ 15 cells per condition. (G) DIC images and cell outlines of spreading MEFs treated with 0.1% DMSO or 10 μM HC-067047. (H-I) Analysis of the effects of TRPV4 suppression on lamellipodial protrusion speed (H) DIC images and cell edge montages of migrating MEFs seeded on fibronectin-coated glass surface in control (siCtrl), TRPV4 depletion (siTRPV4), and rescue conditions. (I) Bar graph of the average protrusion speed of migrating cells in control (siCtrl and DMSO), TRPV4 depletion (siTRPV4), rescue, and TRPV4 inhibition (TRPV4 inh.) conditions. n ≥ 20 cells per condition. Mann-Whitney test is used for comparisons between 2 treatments and one-way ANOVA followed by Tukey-Kramer post-hoc test for comparisons between 3 or more treatments. The data are presented as mean ± standard deviation (SD) unless otherwise specified. * p < 0.05, ** p < 0.01, *** p < 0.005, **** p < 0.001. See also Figures S1, S2, and Video S1.

The concentration of cytoplasmic Ca^2+^ is controlled by the influx of extracellular Ca^2+^ as well as the efflux of intracellular Ca^2+^ from intracellular stores ^28–30^. Despite being the primary storage site for intracellular Ca^2+^, the endoplasmic reticulum (ER) is mainly absent from the cell periphery^31,32^ and is therefore unlikely to be involved in the regulation of local Ca^2+^ level in lamellipodia. To test this directly, we treated cells with thapsigargin, an inhibitor of ER Ca^2+^ ATPases (SERCA pumps)^29^. Disruption of ER Ca^2+^ stores did not alter the GCaMP signal gradient and cell spreading speed compared to control cells (Figures 1D, E and S1F). Therefore, we focused on the contribution of extracellular Ca^2+^ to lamellipodia protrusion. We found that chelation of extracellular Ca^2+^ or suppression of stretch-activated channels on the plasma membrane with GdCl_3_ significantly reduced cell spreading speed (Figures 1D and E), suggesting that influx of extracellular Ca^2+^ through stretch-activated channels is primarily responsible for the control of lamellipodia protrusion.

Given that several members of the stretch-activated transient receptor potential (TRP) channels have been implicated in the local regulation of Ca^2+^ signaling and cell migration^20,33,34^, we next conducted short interfering RNA (siRNA) knockdown experiments targeting Ca^2+^-permeable TPR channels to identify those that might contribute to lamellipodia protrusion. We altered the expression of 18 TRP channels, expressed by fibroblasts (Figure S1G), by using siRNAs and measured cell spreading speed as described above. Depletion of four channels (TRPV1, TRPV4, TRPV5 and TRPV6) suppressed lamellipodia protrusion in spreading cells with TRPV4 having the strongest effect (Figures 1F and G).

Previous work has shown that TRPV4 controls cell-ECM attachment by modulating β1 integrin expression^35^. To examine whether the effect of TRPV4 on lamellipodia is mediated by integrin-based adhesion complexes, we assessed spreading kinetics of control and TRPV4-depleted cells on poly-L-lysine, a substrate that permits integrin-independent cell attachment^36^. These experiments revealed that the reduction in lamellipodia protrusion due to TRPV4 depletion was not influenced by integrin-based adhesion, as evidenced by the slower spreading of TRPV4-depleted cells on poly-L-lysine (Figures S1H and I).

To test the role of extracellular Ca^2+^ influx through TRPV4, we pharmacologically suppressed its channel activity with HC-067047, a selective TRPV4 inhibitor that alters channel permeability by creating a hydrophobic seal in the pore^37^. Cells treated with 10 μM HC-067047 exhibited a significant reduction in the rate of cell spreading (Figures 1F and G). Such changes were conserved across different cell types, including osteosarcoma cells (U2OS), cervical cancer cells (HeLa), and triple-negative breast cancer cells (MDA-MB-231), all of which showed a significant reduction in cell spreading speed when TRPV4 was inhibited (Figures S2A-C). Collectively, these data demonstrate a key role for TRPV4-mediated Ca^2+^ influx in cell spreading and suggest TRPV4 involvement in the control of lamellipodia protrusion.

The changes in cell spreading speed motivated us to examine the role of TRPV4 in lamellipodia protrusion in migrating MEFs. Using immunofluorescence (IF) microscopy, we mapped actin and the ArpC2 subunit of the Arp2/3 complex in control and TRPV4-suppressed MEFs. Among four TRPV4-specific siRNAs, two efficiently reduced TRPV4 expression (Figures S2D-F). The suppression of TRPV4 with siRNAs or HC-067047 significantly decreased the average length of lamellipodia, resulting in ≈40% reduction in the fraction of cell edge positive for ArpC2 (Figures S2G and H). These data indicate that, while not being essential, TRPV4 potentiates lamellipodia formation. Next, we investigated the dynamic behavior of lamellipodia using live-cell differential interference contrast imaging. Consistent with our findings with spreading cells, suppression of TRPV4 led to a ≈25% reduction in protrusion speed in MEFs undergoing random migration, and this effect was fully rescued upon overexpression of siRNA-resistant TRPV4 (Figures 1H and I; Video S1). Collectively, these data identify the mechano-gated Ca^2+^-permeable channel TRPV4 as a key component of a Ca^2+^-dependent signaling pathway regulating protrusion of lamellipodia.

### Local regulation of Ca^2+^ via TRPV4 drives lamellipodia protrusion

To further investigate the role of the TRPV4 channel in the control of Ca^2+^ and protrusion dynamics, we first visualized TRPV4 and F-actin by IF microscopy. Similar to other mechano-gated ion channels^38^, TRPV4 exhibited a uniform distribution across the plasma membrane with a notable portion of TRPV4 localized at lamellipodia (Figure S2I). Inspired by this localization, we next tested whether TRPV4 controls the local Ca^2+^ concentration in lamellipodia. We dynamically mapped the Ca^2+^ biosensor GCaMP6f, soluble mCherry, and F-tractin-emiRFP670 in both control and TRPV4-inhibited MEFs (Figure 2A). Ratiometric imaging revealed a significantly higher GCaMP signal in the protruding cell edge of control cells. TRPV4 inhibition reduced the lamellipodial GCaMP signal by ≈20% within a matter of seconds (Figures 2B-D; Video S2). The suppression of TRPV4 had a negligible effect on the GCaMP signal calculated for the whole cell (Figures 2A and B), suggesting that TRPV4 serves as a local regulator of Ca^2+^ influx in lamellipodia.

**Figure 2.**
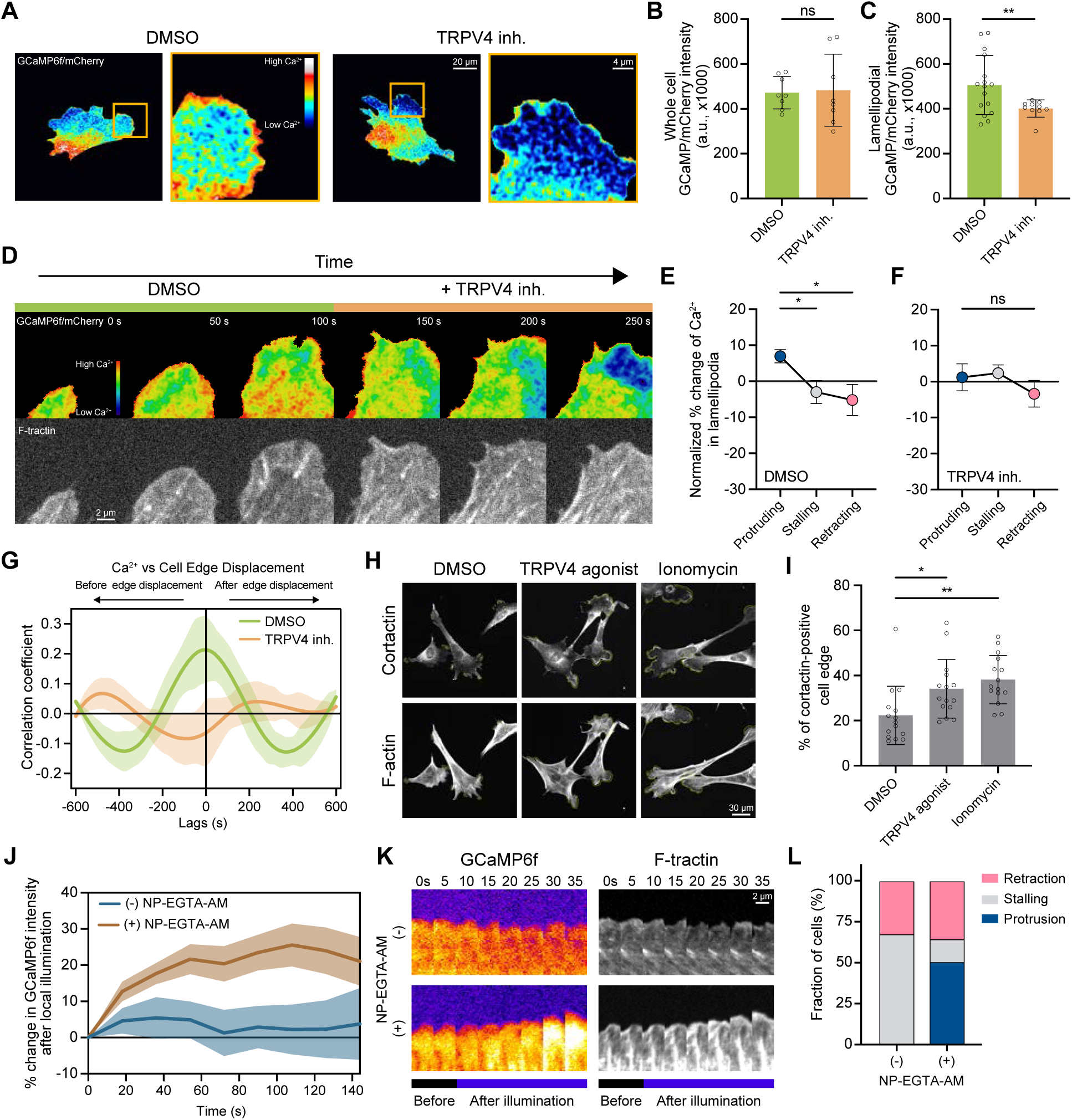
TRPV4 facilitates lamellipodial protrusions by locally controlling Ca^2+^ influx. (A-C) Analysis of intracellular Ca^2+^ levels upon TRPV4 inhibition by ratiometric GCaMP6f/mCherry imaging (A) Ratiometric GCaMP6f/mCherry images of MEFs in 0.1% DMSO and TRPV4 inhibitor (10 μM HC-067047) conditions. (B-C) Quantifications of the (B) whole cell and (C) lamellipodial GCaMP6f/mCherry intensities in the presence of 0.1% DMSO or TRPV4 inhibitor (10 μM HC-067047) conditions. n ≥ 8 cells per condition. (D-F) Analysis of lamellipodial Ca^2+^ level upon acute TRPV4 inhibition (D) Ratiometric GCaMP6f/mCherry and F-tractin-emiRFP670 images of a protruding cell edge before and after TRPV4 inhibition. (E-F) Quantification of the percent change of lamellipodial GCaMP6f/mCherry intensity in protruding (blue), quiescent (dark grey), and retracting (red) phases of lamellipodia. The colored dot indicates the average value from multiple cells. Each data point is normalized to the average intensity during protrusion, stalling and retracting lamellipodia extracted from the same kymograph. n = 7 cells per condition. (G) Cross-correlation analysis between cell edge displacement and lamellipodial Ca^2+^ level. The solid lines indicate the average cross-correlation values of multiple cells. n ≥ 9 cells per condition. The shaded area shows the SE. (H-I) Analysis of the effects of TRPV4 activation on the formation of lamellipodia (H) F-actin and anti-cortactin images of cells treated with 0.1% DMSO, TRPV4 agonist (15 nM GSK1016790A), or 1 μM ionomycin for 5 mins. (I) Quantification of percent of cortactin-positive cell edge in cells treated with 0.1% DMSO, TRPV4 agonist (15 nM GSK1016790A), or 1 μM ionomycin for 5 mins. n = 15 cells per condition. (J-L) Analysis of the effect of photo-released Ca^2+^ on lamellipodia. (J) Quantification of percent change in GCaMP6f intensity after illumination of cells without (-) or with (+) NP-EGTA-AM. n ≥ 10 cells per condition. (K) GCaMP6f and F-tractin-emiRFP670 images of cells before and after illumination without (-) or with (+) NP-EGTA-AM. (L) Responses of cell edge after illumination without (-) or with (+) NP-EGTA-AM. 1 µM ML-7 (a myosin light chain kinase inhibitor). The inhibition of myosin light chain kinase was applied to avoid cell retraction caused by Ca^2+^-dependent myosin activation in the lamella^21^. n ≥ 6 cells per condition. Mann-Whitney test is used for comparisons between 2 treatments and one-way ANOVA followed by Tukey-Kramer post-hoc test for comparisons between 3 or more treatments. The data are presented as mean ± SD unless otherwise specified. * p < 0.05, ** p < 0.01, *** p < 0.005, **** p < 0.001. See also Videos S2 and S3.

Lamellipodia are highly dynamic cellular structures exhibiting rapid cycles of protrusion and retraction^22^; therefore, we tested whether the TRPV4 channel is differentially activated during any distinct phase of the cycle. To test this, we compared the GCaMP signal intensity in protruding, stalling, and retracting lamellipodia of control and TRPV4-inhibited cells (Figures 2D-F). These experiments revealed that in control cells, the Ca^2+^ level was highest in actively protruding lamellipodia, while the transitions to stalling and retracting states coincided with a ≈15% decrease in GCaMP signal intensity. In contrast, the suppression of TRPV4 abolished the pattern of Ca^2+^ influx during the protrusion-retraction cycle, causing the protruding cell edge to stall or collapse (Figure 2D). These data suggest a tight coupling between TRPV4 activation and the protrusion phase of lamellipodia. These findings were further corroborated by a cross-correlation analysis, which revealed a strong positive correlation between Ca^2+^ influx and the onset of lamellipodia protrusion in control, but not TRPV4-inhibited cells (Figure 2G), indicating a key role for TRPV4-mediated Ca^2+^ influx in lamellipodia protrusion.

Next, we sought to examine the effects of an increase in Ca^2+^ influx via TRPV4 on cellular protrusions. We treated MEFs with a TRPV4 agonist (GSK1016790A) and quantified the fraction of the cell edge positive for the lamellipodia marker cortactin.

Ectopic activation of TRPV4 nearly doubled the amount of lamellipodia in fibroblasts (Figure 2H). A comparable increase in lamellipodia formation was observed upon upregulation of Ca^2+^ influx by ionomycin treatment (Figures 2H and I), suggesting that an increase in intracellular Ca^2+^is sufficient to drive protrusion formation. To directly test whether a local increase in intracellular Ca^2+^ is sufficient to induce lamellipodia protrusion, we examined the effects of Ca^2+^ uncaging from the photolabile Ca^2+^ chelator NP-EGTA-AM on the dynamics of the stalling cell edge. Localized uncaging was induced by illuminating a small (3×3 µm) region of the cytoplasm directly adjacent to the stalling cell edge with a low-intensity 405 nm laser. This treatment, while having no noticeable effect on control cells, resulted in a ≈20% increase in GCaMP signal in the vicinity of the illuminated area in NP-EGTA-AM-loaded cells (Figure 2J). Notably, the magnitude of uncaging was quantitatively similar to the Ca^2+^ influx mediated by TRPV4 (Figure 2B). The light-stimulated Ca^2+^ uncaging triggered the protrusion of the stalling cell edge, with over 80% of the tested cells exhibiting a robust protrusion (Figures 2K and L; Video S3), demonstrating that a local increase in intracellular Ca^2+^ is sufficient to induce lamellipodia protrusion. Together, these results indicate that TRPV4 facilitates lamellipodia protrusion through the local control of Ca^2+^ influx.

### TRPV4-mediated Ca^2+^ influx modulates cell edge protrusions through the regulation of F-actin density in lamellipodia

Previous studies have shown that Ca^2+^ influxes control lamellipodia protrusion by triggering myosin activation at distant locations, *i.e.*, within the lamellae, resulting in a partial retraction of the nearby protruding lamellipodia^21^. Therefore, we examined whether the modulation of lamellipodia protrusion by TRPV4 is contingent upon myosin activation. Initially, we utilized IF microscopy to visualize the distribution of the phosphoserine-19 myosin regulatory light chain-2 (pS19MLC2), a marker of myosin activation and contractility^39^, in control and TRPV4-inhibited cells. Consistent with previous reports, these experiments revealed a monotonic gradient of pS19MLC2 intensity, with minimal amounts of pS19MLC2 near the cell edge (Figures S3A and B). This distribution of pS19MLC2 was unaffected by TRPV4 inhibition, indicating that myosin activity is not governed by TRPV4-mediated Ca^2+^ influx.

To directly assess the role of myosin contractility in the regulation of lamellipodia protrusion by TRPV4, we examined the effect of TRPV4 inhibition on cells in which myosin activity was inhibited by blebbistatin (Figures S3C and D). This experiment revealed that suppression of myosin had a minimal effect on the cellular response to TRPV4 inhibition, as the suppression of TRPV4 continued to inhibit lamellipodia protrusion. Collectively, these findings demonstrate that the control of lamellipodia protrusion by TRPV4 is independent of distant changes in myosin contractility, and instead strongly suggest a local Ca^2+^-dependent regulation of the lamellipodial actin network.

We next analyzed the role of TRPV4 in actin organization in lamellipodia. By using IF microscopy, we visualized F-actin and cortactin in control and TRPV4-suppressed cells. We found a significant decrease in lamellipodial actin intensity, with a reduction of 47% in TRPV4-depleted cells and 34% in TRPV4-inhibited cells compared to their respective controls (Figures 3A and B). This marked decline in lamellipodial F-actin coincided with a ≈40% decrease in the intensities of the actin nucleating factor Arp2/3 and cortactin (Figures 3A and C-E). Notably, the recruitment of Abi1, an essential component of the WAVE complex and an upstream regulator of Arp2/3^40^, to lamellipodia was unaffected by TRPV4 inhibition (Figures S3E and F), suggesting that Arp2/3 activity is preserved upon TRPV4 inhibition. Nevertheless, the reduced actin and Arp2/3 staining strongly indicates that suppression of TRPV4 perturbs the organization of the actin network in lamellipodia.

**Figure 3.**
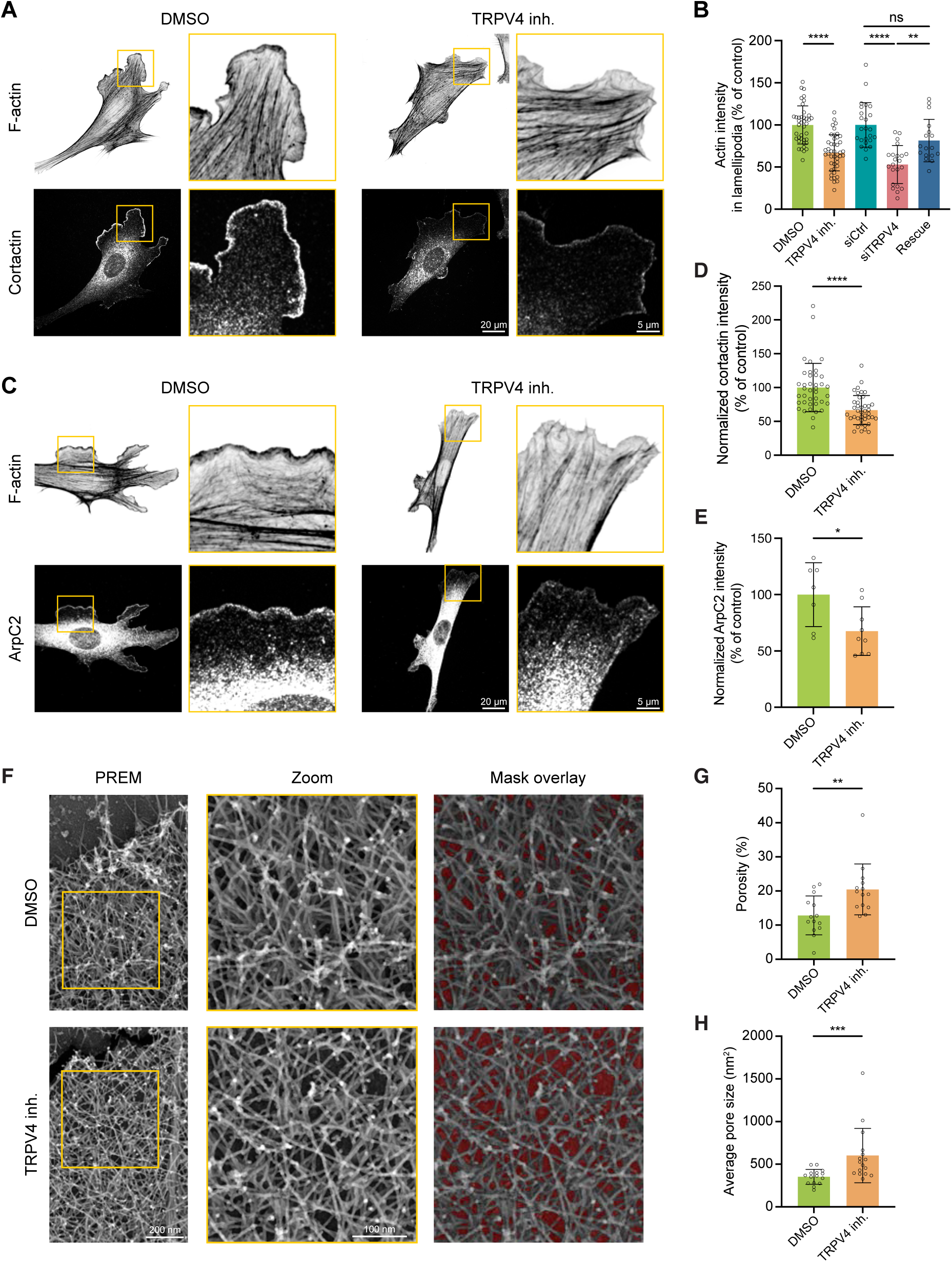
TRPV4-mediated Ca^2+^ influx modulates F-actin density in lamellipodia. (A-E) Analysis of lamellipodial F-actin and lamellipodia markers in MEFs. (A) Fluorescence staining with phalloidin and anti-cortactin of MEFs treated with DMSO or TRPV4 inhibitor. (B) Quantifications of the F-actin intensity in lamellipodia in control conditions (DMSO and siCtrl) and after TRPV4 inhibition or depletion. F-actin intensity of each cell is normalized to the average intensity of control cells. n ≥ 15 cells per condition. (C) Fluorescent phalloidin and anti-ArpC2 images of cells treated with DMSO and TRPV4 inhibitor. Quantifications of (D) lamellipodial cortactin intensity (n > 30 cells per condition) and (E) lamellipodial ArpC2 intensity upon TRPV4 inhibition (n ≥ 7 cells per condition). Each cell’s cortactin or ArpC2 intensity is divided by the average of its control to get a normalized intensity. (F-H) Analysis of lamellipodia F-actin architecture in MEFs using platinum replica electron microscopy (PREM). (F) PREM (left and middle) and the mask overlay images (right) of lamellipodia from MEFs treated with DMSO (top) or TRPV4 inhibitor (bottom). The red areas indicate the voids between actin filaments. Quantifications of lamellipodial F-actin meshwork porosity (G) and the average pore sizes (H) in the presence of DMSO or TRPV4 inhibitor. n ≥ 13 cells per condition. Mann-Whitney test is used for comparisons between 2 treatments and one-way ANOVA followed by Tukey-Kramer post-hoc test for comparisons between 3 or more treatments. The data are presented as mean ± SD unless otherwise specified. * p < 0.05, ** p < 0.01, *** p < 0.005, **** p < 0.001. See also Figure S3.

To directly assess the effects of TRPV4 inhibition on the architecture of lamellipodial actin, we examined F-actin density in cells treated with either DMSO or TRPV4 inhibitor by using platinum replica electron microscopy (PREM). The PREM images did not reveal any gross differences in the organization of the actin network in lamellipodia of control and TRPV4-inhibited cells. Nevertheless, the quantitative analysis of the PREM images revealed ≈60% increase in the sample area of voids between actin filaments, indicating a significant decrease in F-actin density in lamellipodia. Collectively, these findings indicate that TRPV4 facilitates the assembly of the dense network of lamellipodial actin filaments.

### TRPV4 upregulates lamellipodia F-actin density through the RhoA/formin signaling module

We next sought to dissect the molecular mechanism through which TRPV4 regulates the density of F-actin in lamellipodia. Given that the control of actin polymerization is primarily orchestrated by the Rho-family small GTPases, RhoA, Rac1, and Cdc42, we hypothesized that TRPV4 facilitates the assembly of lamellipodial F-actin by activating one or more of these GTPases. To test this, we examined the effect of TRPV4 suppression on the spatiotemporal pattern of RhoA, Rac1, and Cdc42 by using Förster Resonance Energy Transfer (FRET) biosensors. The biosensors were validated by measuring the FRET efficiency of the dominant-negative (RhoaA^T19N^, Rac1^T17N^, and Cdc42^T17N^) and constitutively active (RhoA^Q63L^, Rac1^Q61L^, and Cdc42^Q61L^) mutants (Figures S4A-E). In agreement with previous reports^11,41,42^, FRET imaging revealed a broad gradient of Rac1 and Cdc42 activity across the protruding edges of control cells with the highest activity localized at the tip of the cell edge (Figures S4B and C). Notably, neither the integrated FRET efficiency (FRET_eff_) signal nor the distribution of Rac1 and Cdc42 FRET intensity across the cell edge were significantly affected by the suppression of TRPV4 (Figures S4B-E). In agreement with these data, activity-based Rac1- and Cdc42-GTP pull-down assays did not reveal a significant change in the average cellular activity of Rac1 and Cdc42 upon TRPV4 inhibition (Figures S4F and G). Together, these data demonstrate that TRPV4 is unlikely to regulate lamellipodial actin through the modulation of Rac1 and Cdc42 activity.

Previous studies have demonstrated the activation of RhoA at protruding lamellipodia and established that it plays a role in the assembly of actin filaments that are competent for Arp2/3 binding^11,17,43^. Hence, we initially assessed RhoA activity in control and TRPV4-inhibited cells using an affinity-based pull-down assay. We found that suppression of TRPV4 had a minimal impact on the overall cellular activity of RhoA (Figure 4A), mirroring the unchanged myosin contractility upon TRPV4 inhibition (Figures S3A and B). To determine if the spatial pattern of RhoA activity is altered upon TRPV4 suppression, we compared the distribution of active RhoA, visualized by a FRET biosensor, in control and TRPV4-inhibited cells. Consistent with previous reports^11,44^, RhoA activity exhibited a peak at the tips of control cell protruding edges and gradually decreased towards the cell body (Figures 4B-D). However, TRPV4-inhibited cells exhibited a distinct spatial pattern of RhoA activity, despite maintaining a comparable overall activity level (Figures 4B-E). In TRPV4-inhibited cells, active RhoA was no longer enriched in the lamellipodia, as evidenced by a flattened intensity profile and a reduced lamellipodia-to-cell body FRET_eff_ ratio (Figures 4D and F), suggesting that TRPV4-mediated Ca^2+^ influx activates RhoA at protruding cell edges.

**Figure 4.**
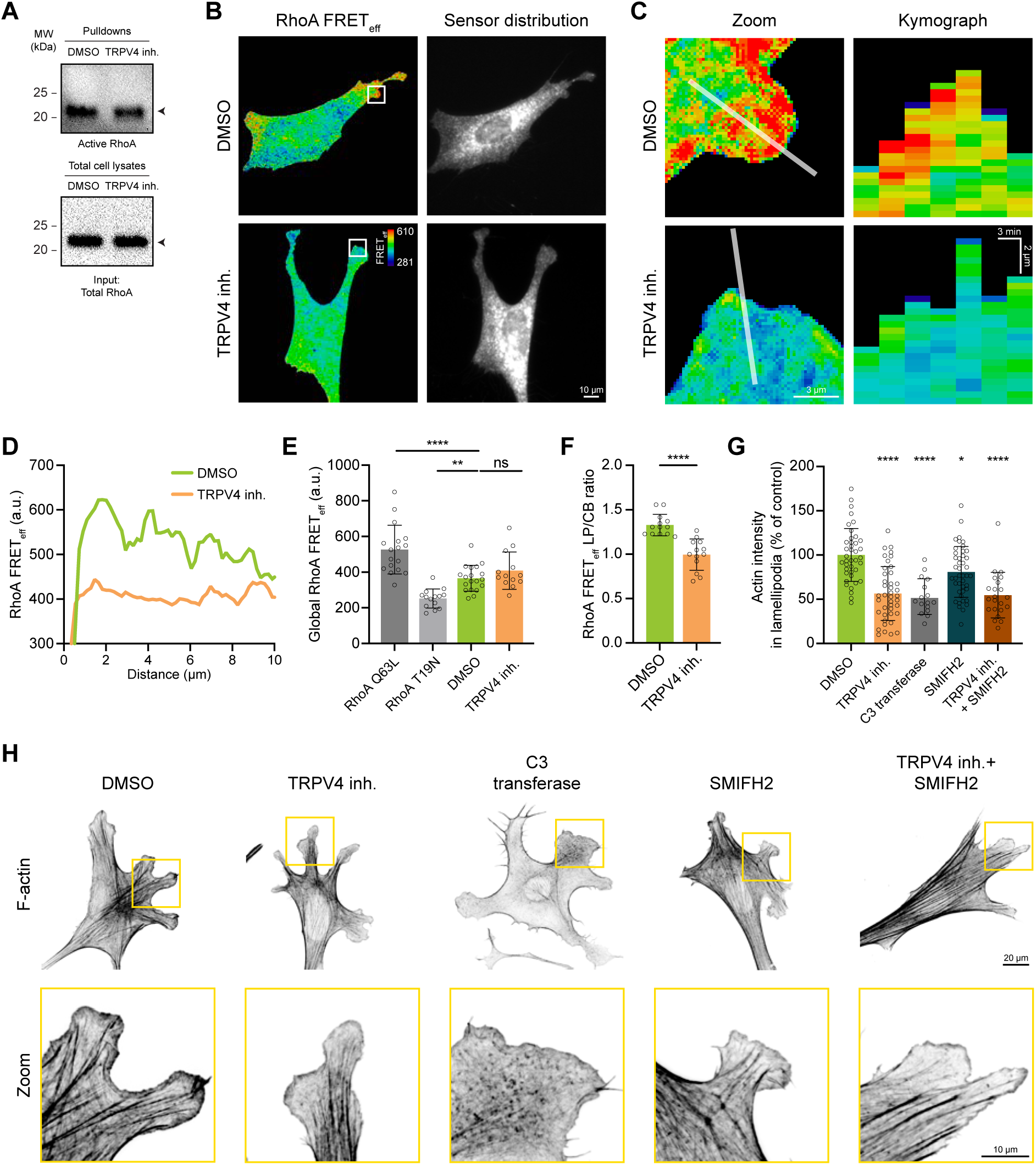
TRPV4 upregulates lamellipodial F-actin density via the RhoA/formins pathway. (A) Western blot showing the activity of RhoA in whole cell lysates treated with 0.1% DMSO or TRPV4 inhibitor (10 μM HC-067047) by using the active GTPase pull-down assay. (B-F) RhoA FRET analysis of control and TRPV4-inhibited cells. (B) Fluorescence images of RhoA FRET biosensor activation and its distribution (acceptor channel). The white box indicates the cell edge region used for kymograph analysis. (C) (Left) Zoomed RhoA FRET images of the cell edge in the DMSO and TRPV4-inhibited cells. The white line indicates the region used to create the kymograph on the right. (Right) Kymographs of protruding cell edges in DMSO and TRPV4 inhibitor. (D) Line intensity profiles of RhoA FRETeff across cell edges in DMSO and TRPV4 inhibitor. (E) Quantifications of global RhoA FRETeff. (F) Quantifications of lamellipodia (LP)-to-cell body (CB) RhoA FRETeff. n ≥ 10 cells per condition. (G-H) Effects of suppressing the RhoA/formins signaling pathway on lamellipodial F-actin. (G) Quantifications of lamellipodial F-actin intensity in different conditions. n > 15 cells per condition. (H) Fluorescent phalloidin staining images of cells treated with 0.1% DMSO, TRPV4 inhibitor (10 μM HC-067047), Rho inhibitor (C3 transferase, 2.0 µg/ml, 3 hr), pan-formins inhibitor (SMIFH2, 10 µM, which has a negligible effect on myosin-II activity^83^), or a combination of TRPV4 and formin inhibitors. Mann-Whitney test is used for comparisons between 2 treatments and one-way ANOVA followed by Tukey-Kramer post-hoc test for comparisons between 3 or more treatments. The data are presented as mean ± SD unless otherwise specified. * p < 0.05, ** p < 0.01, *** p < 0.005, **** p < 0.001. See also Figure S4.

Next, we sought to determine whether the activation of RhoA/formin(s) signaling module is sufficient to explain the effect of TRPV4 inhibition on the actin density in lamellipodia. To test this, we examined actin density in cells treated with TRPV4 or RhoA inhibitors (C3 transferase) and immunostained with cortactin antibody and phalloidin.

In agreement with previous reports^45,46^, cells treated with C3 transferase exhibited a significant reduction in the number and intensity of stress fibers (Figures 4H), indicating a decline in Rho-mediated myosin contractility. Conversely, TRPV4 inhibition had a minimal effect on stress fibers, in line with our finding of no significant change in the overall cellular RhoA activity upon TRPV4 inhibition. However, cells treated with either TRPV4 inhibitor or C3 transferase exhibited a comparable (≈40%) decrease in the intensity of the lamellipodia network (Figures 4G and H), supporting the notion of TRPV4- and RhoA-dependent regulation of actin in lamellipodia. Furthermore, suppression of the downstream RhoA effectors, formins, partially recapitulated the phenotype of TRPV4 inhibited cells as evident from the 20% decrease in lamellipodia actin density (Figures 4G and H). Co-inhibition of formins and TRPV4 did not additively decrease actin density, suggesting that lamellipodia actin is controlled through the TRPV4-dependent activation of the RhoA/formin(s) signaling module, rather than by the convergence of these pathways on actin regulation. Together, these data identify a new signaling module composed of TRPV4/RhoA/formins that control the organization of lamellipodia actin network.

### CaMKII facilitates lamellipodia protrusion and upregulates RhoA activity in a TRPV4-dependent manner

We next sought to examine whether the control of lamellipodia architecture and protrusion dynamics by TRPV4 is mediated by Ca^2+^/calmodulin-dependent protein kinase II (CaMKII), which is known to remodel the actin cytoskeleton through small GTPases in neuronal spines^47^. In non-neuronal cells, including vascular smooth muscle cells and cancer cells, CaMKII has been implicated in migration and invasion^48,49^.

We first examined the level of active CaMKII (pCaMKII^T286^) in control and TRPV4-inhibited cells by using western blotting and immunostaining with a phospho-specific antibody. Inhibiting TRPV4, much like its effect on intracellular Ca^2+^ concentration, did not alter overall CaMKII activity, as indicated by similar pCaMKII intensities in lysates of control and TRPV4-inhibited cells (Figure S5A). However, IF microscopy revealed a significantly lower level of pCaMKII in the lamellipodia of TRPV4-inhibited cells, accompanied by reduced F-actin intensity (Figures 5A-C). Moreover, quantitative analysis revealed a strong positive correlation between local CaMKII activity and actin density in lamellipodia (Figure 5C), suggesting that CaMKII is involved in the regulation of the lamellipodial actin network.

**Figure 5.**
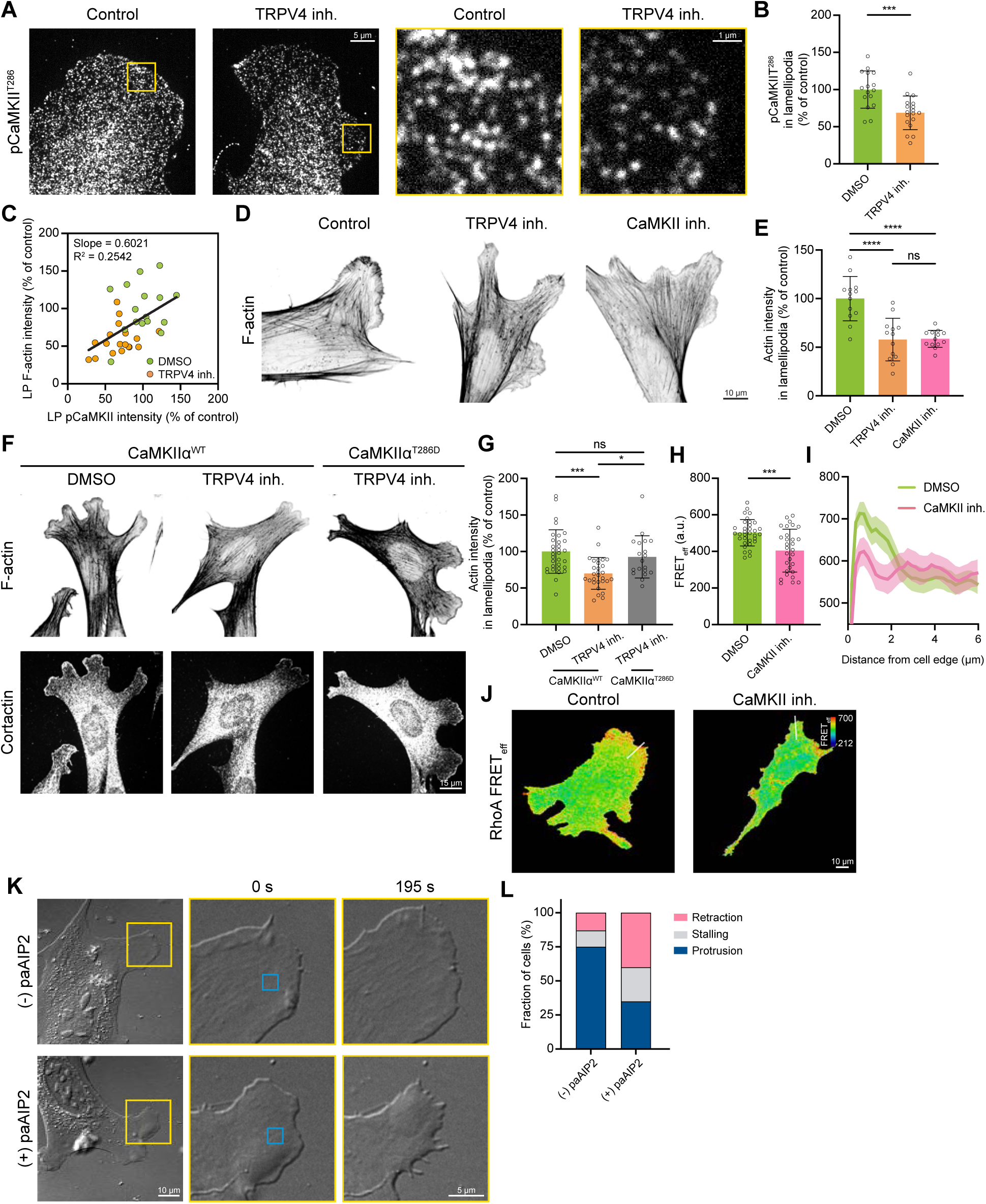
TRPV4-regulated CaMKII signaling upregulates local RhoA activity and facilitates cell protrusion. (A-C) Analysis of pCaMKII^T286^ level in MEFs treated with either 0.1% DMSO or TRPV4 inhibitor (10 μM HC-067047). (A) Images of anti-pCaMKII^T286^ and fluorescent phalloidin staining. (B) Quantifications of lamellipodial pCaMKII^T286^. n > 15 cells per condition. (C) A scatter plot of lamellipodial (LP) F-actin and pCaMKII intensities of individual cells treated DMSO or TRPV4 inhibitor. The black line indicates the best-fit linear regression. (D-E) Effects of CaMKII inhibition (10 μM KN-93) on F-actin intensity. (D) Fluorescent phalloidin staining images of cells treated with DMSO, TRPV4 inhibitor, or CaMKII inhibitor. (E) Quantifications of lamellipodial F-actin intensity. n = 12 cells per condition. (F-G) Effects of phosphomimetic CaMKII (CaMKIIα^T286D^) on F-actin in the presence of TRPV4 inhibitor. (F) Fluorescent phalloidin and anti-cortactin staining images in cells expressing wildtype CaMKIIα in DMSO or TRPV4 inhibitor, or CaMKIIα^T286D^ in TRPV4 inhibitor. (G) Quantifications of their LP F-actin intensity. n > 15 cells per condition. (H-J) Effects of CaMKII inhibition on RhoA activity. (H) Quantifications of global RhoA FRETeff in cells treated with DMSO or CaMKII inhibitor (n > 15 cells per condition) and (I) the average RhoA FRET intensity profiles extracted near the cell edges of cells treated with DMSO or CaMKII inhibitor. n = 15 cells per condition. The shaded area shows the standard error. The locations where the intensity profiles are taken are indicated in the corresponding FRET images with white lines. (J) RhoA FRET images of MEFs treated with DMSO or CaMKII inhibitor. (K-L) Optogenetic inhibition of CaMKII in protruding cell edges using paAIP2. (K) DIC images of cells without (-) or with (+) paAIP2. The yellow box indicates the cell edge region used for the zoomed-in images shown on the right. The indicated time shows the time elapsed after the light stimulation. The blue box indicates the region of light stimulation (405 nm) region. (L) Fractions of cells undergoing protrusion, stalling, or retraction post light stimulation. n ≥ 8 cells per condition. Mann-Whitney test is used for comparisons between 2 treatments and one-way ANOVA followed by Tukey-Kramer post-hoc test for comparisons between 3 or more treatments. The data are presented as mean ± SD unless otherwise specified. * p < 0.05, ** p < 0.01, *** p < 0.005, **** p < 0.001. See also Figure S5 and Video S4.

Next, we sought to determine whether actin density in lamellipodia is regulated by CaMKII downstream of TRPV4. To test this, we compared the intensities of phalloidin-stained actin in lamellipodia of cells treated with CaMKII or TRPV4 inhibitors. We found that suppression of CaMKII with 10 μM KN-93 reduced F-actin intensity by 42%, phenocopying the effect of TRPV4 inhibition (Figures 5D and E). Furthermore, the effect of TRPV4 inhibitor was fully rescued by ectopic activation of CaMKII, as evidenced by a significant increase in lamellipodial actin intensity in TRPV4-inhibited cells expressing a constitutively active CaMKII mutant (CaMKIIα^T286D^)^49^, compared to the cells expressing wild-type CaMKIIα (Figures 5F and G). Collectively, these data strongly suggest that CaMKII acts as a downstream effector of TRPV4, enabling the Ca^2+^-dependent regulation of F-actin in lamellipodia.

To determine whether CaMKII regulates RhoA, we assessed the effect of CaMKII inhibition with 10 μM KN-93 on the activity of RhoA by using FRET biosensor. We found that CaMKII suppression led to a modest (≈20%) decrease in RhoA activity throughout the cell (Figure 5H). These experiments also showed a reduced RhoA activity in protrusions of CaMKII inhibited cells (Figures 5I and J), identifying CaMKII as an upstream regulator of RhoA in lamellipodia and suggesting its involvement in the control of protrusive activity. To assess the functional role of CaMKII activation downstream of TRPV4 in lamellipodia protrusions, we employed an optogenetic inhibitor of CaMKII (paAIP2)^50^. Local suppression of CaMKII activity in protruding lamellipodia of MEFs using 405 nm laser stimulation resulted in rapid cessation of protrusion within a few minutes (Figures 5K and S5B; Video S4). Quantification of the protrusion response upon local CaMKII inhibition showed that over 60% of cells switched from protruding to stalling or retracting (Figure 5L). Together, these data establish CaMKII as a downstream effector of TRPV4 that enables Ca^2+^-dependent regulation of lamellipodial actin architecture and protrusion by modulating local RhoA activity.

### TEM4, a RhoGEF, links local activity of TRPV4/CaMKII signaling module to RhoA regulation

Our finding that CaMKII provides a local regulation of lamellipodial RhoA points to the activation of RhoA through phosphorylation. However, the known phosphorylation sites on RhoA, Ser26 or Ser188, have previously been reported to inactivate RhoA upon phosphorylation^51,52^, making it unlikely that CaMKII activates RhoA via direct phosphorylation. Therefore, we hypothesized that TRPV4 regulates RhoA activity in lamellipodia through CaMKII-mediated phosphorylation of Rho GTPase regulators, such as Guanine nucleotide Exchange Factors (GEFs) and/or GTPase-Activating Proteins (GAPs). To identify the RhoA regulators phosphorylated in a CaMKII/TRPV4 dependent manner, we conducted phosphoproteomic profiling of control, TRPV4-inhibited, and CaMKII-inhibited cells. To improve the sensitivity of the assay for low abundance proteins, we biochemically isolated the protrusion fractions as described previously^53,54^.

MEFs cultured on FN-coated polycarbonate membranes with 3 μm pores were exposed to a gradient of chemoattractant (1—10% FBS), which induces protrusions extending through the pores (Figure 6A). This experimental setup effectively triggered protrusions while restricting cell body translocation as evidenced by the positive F-actin staining on both the upper and lower surface of the membrane, and the lack of DAPI staining on the lower surface (Figure 6B). The purity of the isolated protrusion fraction was further confirmed by western blotting against vasodilator-stimulated phosphoprotein (VASP), a molecular marker for the protrusion fraction^55^, as the isolated fractions of cellular protrusions exhibited a significant enrichment with VASP compared to the cell body fraction (Figure S6A).

**Figure 6.**
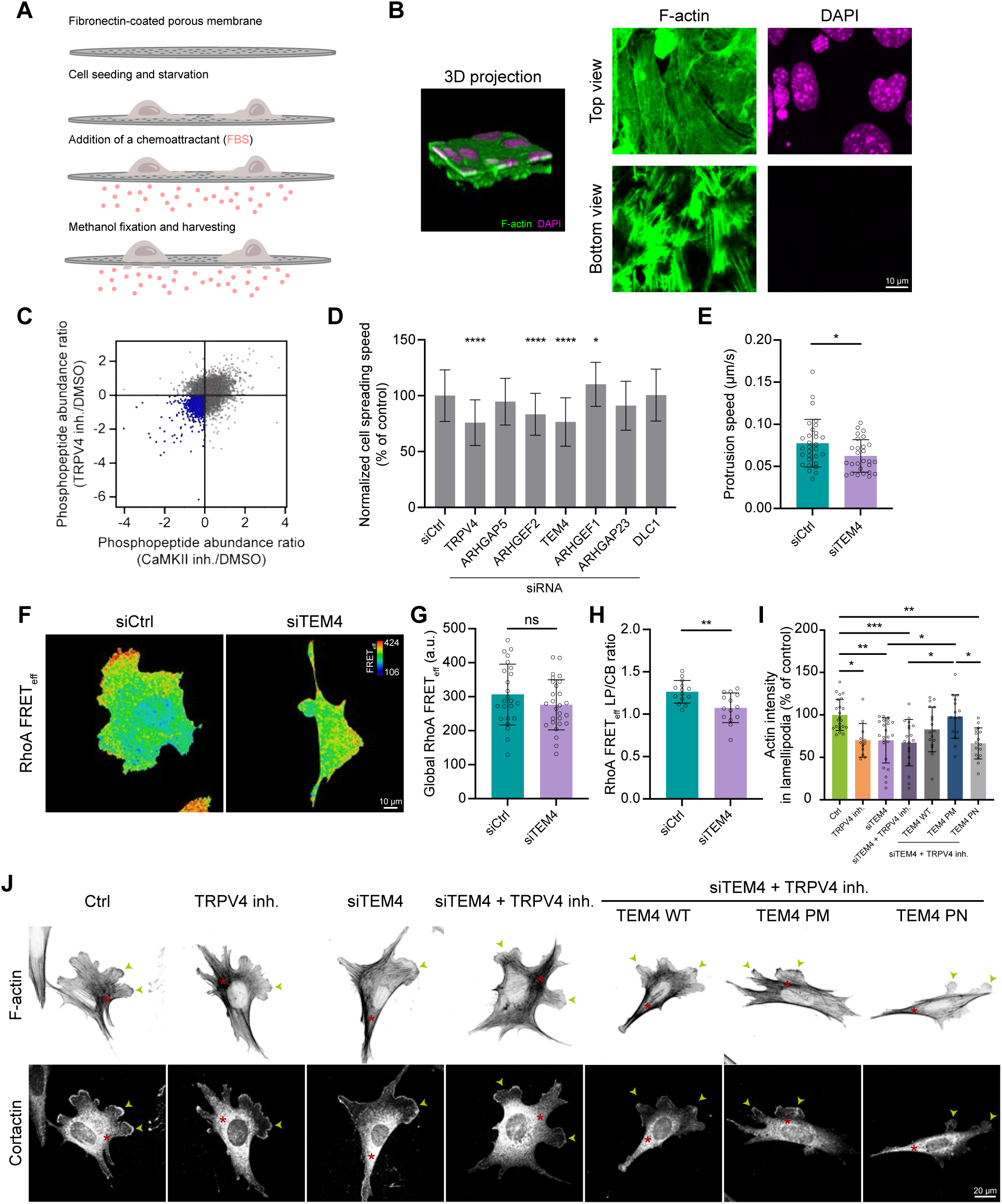
TEM4, a RhoGEF, regulates RhoA activity and lamellipodial protrusions downstream of the TRPV4/CaMKII signaling module. (A-C) Phosphoproteomic analysis of purified lamellipodia (LP) and cell body (CB) fractions. (A) Schematic showing the general protocol of LP and CB fractionation. (B) DAPI and F-actin fluorescence images of MEFs plated on the porous membrane in a transwell. (Left) 3D projection of a 21 µm z-stack acquired at 0.25 µm steps. (Right) Maximum z projected images of the top and bottom compartments. (C) Scatterplot showing the abundance ratio of all the detected phosphopeptides from the phosphoproteomic analysis. The y-axis represents the abundance ratio of TRPV4 inhibition relative to the DMSO control, while the x-axis represents the abundance ratio of CaMKII inhibition relative to the DMSO control. Each dot corresponds to a single phosphopeptide, with blue dots indicating peptides that are commonly down-phosphorylated upon inhibition of TRPV4 and CaMKII. n = 3 biological repeats. (D) Quantifications of the cell spreading speed (relative to the mean of control) upon the siRNA depletion of different RhoGEFs and RhoGAPs. n ≥ 45 cells per condition. (E) Quantifications of the protrusion speed of migrating MEFs in the control and upon TEM4 depletion. n > 25 cells per condition. (F-H) RhoA FRET analysis upon TEM4 depletion. (F) Fluorescence images of RhoA FRET biosensor activation in control (siCtrl) and upon TEM4 depletion. (G) Quantifications of global RhoA FRETeff. n > 20 cells per condition. (H) Quantifications of lamellipodia (LP)-to-cell body (CB) RhoA FRETeff. n ≥ 14 cells per condition. (I-J) Analysis of lamellipodial F-actin intensity in TEM4 phospho-mutant expressing MEFs. (I) Quantifications of the F-actin intensity in lamellipodia of control (DMSO), TRPV4-inhibited, TEM4-depleted, wild type TEM4 rescue, and TEM4 phospho-mutant expressing MEFs (PN: phospho-null; PM: phospho-mimetic) The mutated serine/threonine sites are S142, S145, T150, and S152. Each cell’s lamellipodial F-actin intensity is divided by the average of its control to get a normalized actin intensity relative to its control. n ≥ 15 cells per condition. (J) Fluorescent phalloidin and anti-cortactin images of MEFs. The green arrowheads and red asterisks indicate lamellipodia and actin stress fibers, respectively. Mann-Whitney test is used for comparisons between 2 treatments and one-way ANOVA followed by Tukey-Kramer post-hoc test for comparisons between 3 or more treatments. The data are presented as mean ± SD unless otherwise specified. * p < 0.05, ** p < 0.01, *** p < 0.005, **** p < 0.001. See also Figure S6.

Phosphoproteomic analysis of lamellipodia fractions isolated from control, TRPV4-inhibited, and CaMKII-inhibited cells (Figure 6C), followed by median MS intensity analysis of three biological replicates per condition, identified 1447 phosphopeptides with reduced phosphorylation level upon TRPV4 and CaMKII inhibition (Table S1).

Following bioinformatic analysis, six RhoA regulators (Arhgap5, Arhgef2, Arhgef17, Arhgef1, Arhgap23, and DLC1) were identified. We then siRNA-depleted each candidate protein and evaluated lamellipodia protrusive activity in a cell spreading assay (Figure 6D). Depletion of Arhgef2 (GEF-H1) and Arhgef17 (Tumour Endothelial Marker 4, TEM4) resulted in a reduced lamellipodia protrusion speed similar to that seen with TRPV4 inhibition.

Previous studies have shown that GEF-H1 regulates cellular protrusions by promoting focal adhesion turnover^56^. However, our experiments demonstrated an integrin-independent role of TRPV4 in regulating protrusion dynamics (Figures S1H and I), suggesting that GEF-H1 is unlikely to mediate TRPV4’s effects on lamellipodia. This was further supported by the negligible effect of GEF-H1 depletion on lamellipodia actin density (Figures S6B and C). Together these data make it unlikely that TRPV4-mediated Ca^2+^ influx promotes lamellipodial protrusion via GEF-H1 and leave TEM4 as the most plausible candidate to link TRPV4/CaMKII activity to RhoA.

To examine the role of TEM4 in the regulation of lamellipodia protrusions, we depleted TEM4 by using smart pool siRNAs (Figures S6D and E) and investigated the dynamic behavior of lamellipodia using live cell imaging. Consistent with our findings on spreading cells, depletion of TEM4 led to a ≈20% reduction in protrusion speed in MEFs undergoing random migration (Figure 6E). Next, we compared the pattern of RhoA activity visualized with the FRET biosensor in control and TEM4-depleted cells. This analysis revealed only a minimal, not statistically significant change in the overall RhoA activity upon TEM4 depletion (Figures 6F and G). However, TEM4-depleted cells exhibited an altered spatial pattern of RhoA activity characterized by a lower level of active RhoA in lamellipodia compared to the control cells (Figure 6H). Similar to the effect of TRPV4 inhibitor, depletion of TEM4 efficiently abolished the enrichment of active RhoA at the protruding cell edge as evident from a lamellipodia-to-cell-body RhoA activity ratio of one (Figures 6F and H). TEM4 depletion also phenocopied the effect of TRPV4 inhibition on the organization of the actin cytoskeleton. While depletion of TEM4 did not affect contractile cytoskeletal structures, *e.g.*, stress fibers and actin arches, it markedly decreased the density of the lamellipodia actin network (Figures 6I and J). Inhibiting TRPV4 in TEM4-depleted cells did not additively decrease lamellipodial actin density, suggesting that the control of TEM4 activity was downstream of TRPV4. Together, these data demonstrate that TEM4 is a key mediator that links TRPV4 signaling to RhoA activation in the regulation of lamellipodia protrusions.

To determine the role of TEM4 phosphoregulation in the control of lamellipodia actin network, we generated phospho-mimetic (TEM4^S142D/S145D/T150D/S152D^) and phospho-null (TEM4^S142A/S145A/T150A/S152A^) mutants of TEM4 by substituting serine/threonine residues phosphorylated in a TRPV4/CaMKII-dependent manner with aspartic acid and alanine, respectively. Notably, bioinformatic analysis has identified serine residues 142, 145 and 152 to be a potent CaMKII-targeting site (Figure S6F). We expressed wildtype TEM4 or its mutants in TEM4-depleted cells and examined their effect on lamellipodia actin density upon TRPV4 inhibition. Expression of wildtype TEM4 and the phospho-null mutant did not significantly affect actin density in TRPV4 inhibited cells compared to control. In contrast, the phospho-mimetic TEM4 mutant fully rescued lamellipodial actin density under TRPV4 inhibition (Figure 6I). Thus, phosphorylation of TEM4 is essential for the assembly of dense actin network in lamellipodia. Together, these data demonstrate that N-terminal phosphorylation of TEM4 by the TRPV4/CaMKII signaling axis controls the organization of lamellipodia actin network.

### The TRPV4 signaling axis facilitates cell protrusion and cell migration in vitro and in vivo

To determine the physiological significance of the signaling events triggered by TRPV4-mediated Ca^2+^ influx, we assessed cell migration in vitro and in vivo under the experimental conditions that suppress the key components of TRPV4/CaMKII/TEM4/RhoA signaling axis. Considering the strong effect of RhoA and CaMKII inhibition on cytoskeletal structures and focal adhesions^57,58^, we focused on cells treated with TRPV4 inhibitor or depleted for TEM4. As shown above (Figures 1H, I and 6I), these treatments efficiently suppressed lamellipodia protrusion, while having a minimal effect on cell morphology (Figures S7A-D). Automated tracking of individual cells revealed that both TRPV4-inhibited and TEM4-depleted cells migrated markedly slower than control cells (Figures 7A, B and D; Video S5). We also calculated the directionality index (DI = Displacement/Path; Figures 7C) from cell motion tracks, where a higher DI indicates more directionally persistent movement^59^. This analysis showed that suppression of TRPV4 or depletion of TEM4 had a negligible effect on cell migration directionality (Figure 7E), indicating that TRPV4 and TEM4 are essential for efficient cell migration but dispensable for the control of directionality.

**Figure 7.**
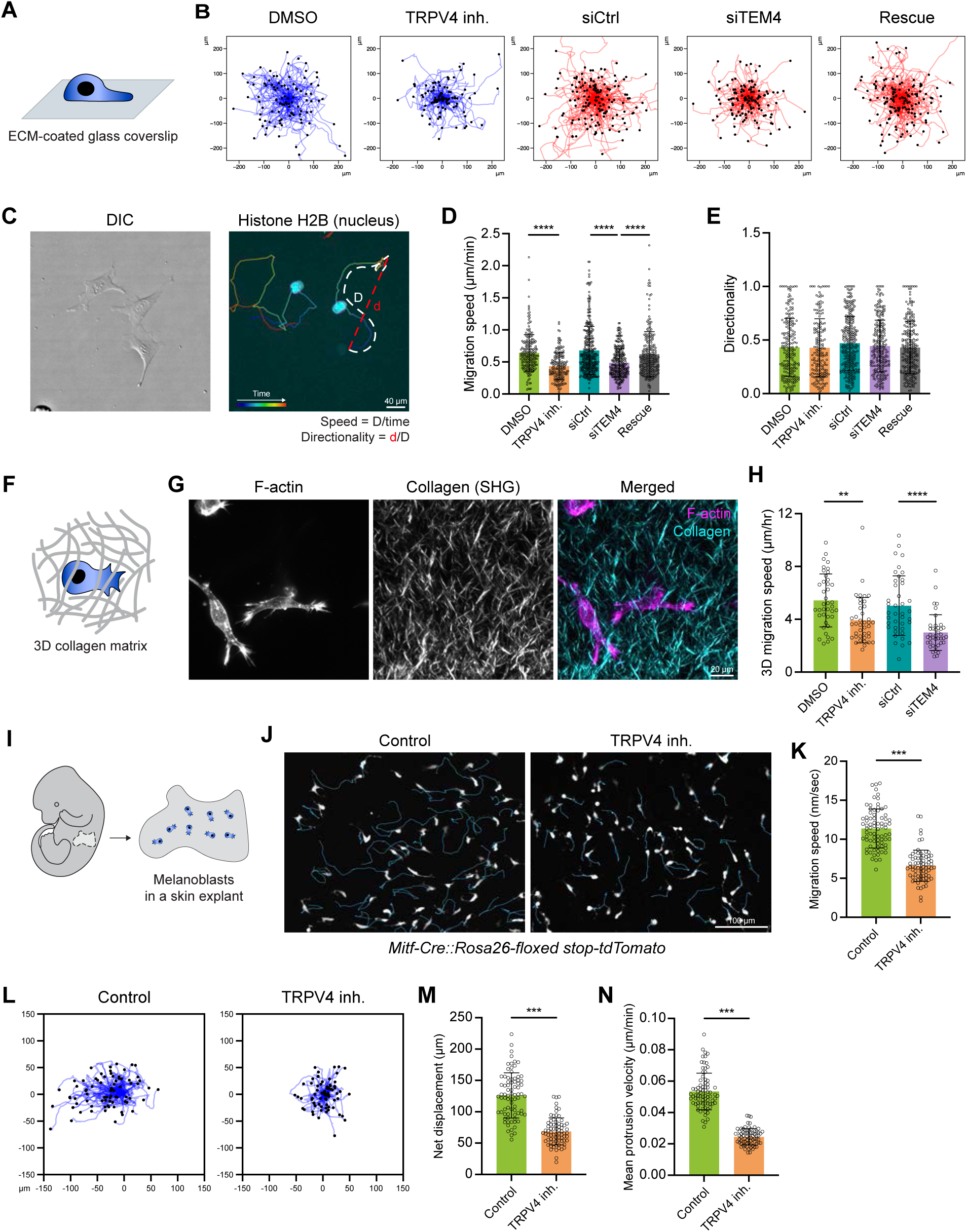
TRPV4/CaMKII/TEM4/RhoA signaling axis regulates cell migration. (A-E) Analysis of the role of TRPV4 signaling axis on random cell migration on fibronectin (FN)-coated substrate in 20 hr. (A) Schematic of the experimental setup for the random migration assay. (B) Migration tracks of MEFs treated with 0.1% DMSO, TRPV4 inhibitor (10 μM HC-067047), non-targeting siRNA (control), or TEM4 siRNA. The black dot indicates the end of the track. Only cell tracks that were continuously tracked for 40 frames were displayed. (C) DIC and fluorescence images of MEFs transiently expressing mEmerald-tagged Histone H2B with the migration tracks overlay. The white and red dotted lines indicate the track length (D) and the Euclidean distance (d), respectively. (D) Quantifications of the average migration speed upon TRPV4 inhibition, and TEM4 depletion and (E) their directionality. n ≥ 182 cells per condition. (F-H) Analysis of the role of TRPV4 signaling axis in 3D migration. (F) Schematic of the experimental setup for the 3D collagen migration assay, in which MEFs stably expressing PH-Akt-eGFP were embedded in the collagen I matrix prior to imaging. (G) Maximum z-projected fluorescent phalloidin staining and second harmonic generation (SHG) images of MEFs embedded in a collagen matrix. (H) Quantifications of average cell migration speed of MEFs over 12 hr stained for Hoechst embedded in collagen matrices. MEFs were treated with TRPV4 inhibition or TEM4 siRNA depletion. n = 40 cells per condition. (I-J) Effects of TRPV4 inhibition on melanoblast migration *ex vivo*. (I) Schematic of the experimental setup for the mouse melanoblast migration assay. (J) Fluorescence images of melanoblasts from *Mitf-Cre::Rosa26-floxed stop-tdTomato* E15.5 embryo trunk skin explant with or without TRPV4 inhibitor. (K) Quantifications of the average melanoblast migration speed in control and upon TRPV4 inhibition. (L) Melanoblast migration tracks in the mouse skin explants. (M-N) Quantifications of net displacement and mean protrusion velocity upon TRPV4 inhibition. n ≥ 74 cells per condition. Mann-Whitney test is used for comparisons between 2 treatments and one-way ANOVA followed by Tukey-Kramer post-hoc test for comparisons between 3 or more treatments. The data are presented as mean ± SD unless otherwise specified. * p < 0.05, ** p < 0.01, *** p < 0.005, **** p < 0.001. See also Figure S7, Videos S5 and S6.

To determine whether the impaired migration observed upon TRPV4 and TEM4 suppression can be attributed to changes in cell-ECM attachment or focal adhesion maturation, we analyzed focal adhesion size in control, TRPV4-inhibited and TEM4-depleted MEFs by using IF microscopy (Figure S7E). TRPV4-inhibited and TEM4-depleted MEFs exhibited focal adhesions comparable in size to those in control cells. The fraction of nascent and mature adhesions was also unaffected, suggesting that TRPV4 and TEM4 suppression does not impact focal adhesion dynamics (Figures S7F-G).

Consistent with the effect of TRPV4 inhibition on cell spreading on poly-L-lysine substrates (Figures S1H and I), these data strongly indicate that the reduced migration observed upon TRPV4 and TEM4 suppression is not due to changes in integrin adhesion complexes.

Previous studies have demonstrated that lamellipodia protrusions facilitate cancer cell migration^60^, therefore we sought to determine the role of the TRPV4 signaling axis in cell migration under physiological conditions. To address this question, we employed two complementary migration assays: collagen scaffolds to assess the morphodynamic properties and migratory behavior of cells in a three-dimensional (3D) environment, and skin explants to examine migration within a native tissue context. To assess the role of TRPV4 and TEM4 in cell migration in collagen scaffolds, we labelled MEFs with a nuclear dye, embedded them in a collagen gel, and visualized cell movement by timelapse spinning disk confocal microscopy (Figures 7F and G). Similar to cells migrating on a two-dimensional (2D) surface, cells in a collagen gel exhibited a significant reduction in cell migration speed upon suppression of TRPV4 and TEM4 (Figure 7H). To determine whether the reduced migration observed in TRPV4- and TEM4-suppressed cells was attributable to impaired lamellipodia, we analyzed the morphology of dendritic protrusions in collagen-embedded cells. We used these structures as a proxy of lamellipodia in 3D environment as their formation requires the assembly of Arp2/3-mediated actin network^60^. We found that suppressing components of the TRPV4 signaling axis led to a reduced length and number of dendritic protrusions (Figure S7H), indicating that the control of lamellipodia protrusive activity by TPRV4 signaling axis is crucial for cell migration, not only in 2D, but also in 3D microenvironments.

Next, we utilized mouse melanoblast migration in a skin explant as a model (Figure 7I). Since the migration of melanoblasts is known to rely on lamellipodial protrusion^9^, we hypothesized that pharmacological inhibition of TRPV4 would suppress lamellipodial protrusion in melanoblasts and consequently impair their migration. To test this hypothesis, we tracked the cell shape and quantified the migration of melanoblasts in the absence or presence of TRPV4 inhibitor. TRPV4 inhibition of melanoblasts in skin explants resulted in a significantly rounder cell morphology compared to the control cells (Figures S7I and J). TRPV4-inhibited melanoblasts exhibited a lower protrusion speed and migrated at a significantly reduced speed compared to the control cells (Figures 7J-N and S7K; Video S6), demonstrating that Ca^2+^ influx through TRPV4 is required for the protrusion and migration of melanoblasts in vivo. Collectively, these data underscore the importance of the TRPV4 signaling axis in regulation of cell migration in vitro and in vivo.

## DISCUSSION

Our study reports the discovery of a novel signaling axis that regulates the protrusive activity of migrating cells by facilitating the assembly of the actin network in lamellipodia. By screening a panel of Ca^2+^-modulating drugs and siRNAs targeting Ca^2+^ channels in a cell spreading assay, we identified a stretch-activated channel, TRPV4, as a positive regulator of lamellipodia across various cell types. Quantitative imaging revealed that TRPV4 suppression reduces Ca^2+^ influx in lamellipodia, which subsequently reduces lamellipodia actin density and protrusive activity. We further demonstrate that the coupling between Ca^2+^ influx and F-actin density is mediated by a Ca^2+^-dependent kinase CaMKII, which phosphorylates a RhoGEF TEM4 resulting in a local activation of RhoA. This activation, in turn, facilitates formin-mediated actin polymerization, driving the assembly of a dense lamellipodial actin network. Disrupting this signaling axis (TRPV4/CaMKII/TEM4/RhoA) impairs lamellipodial actin assembly and diminishes protrusive activity. Importantly, we demonstrate the physiological relevance of this signaling axis in regulating cell migration *in vitro* and *in vivo*.

Previous studies have demonstrated the key role of Ca^2+^ in the regulation of cell migration^29,61^. Influx of extracellular Ca^2+^ through stretch-activated ion channels has been shown to control myosin activity in the cell body, coordinating protruding activity at the leading edge with retraction of the trailing edge^18^. Transient activation of Ca^2+^ channels localized on the plasma membrane and endoplasmic reticulum has also been shown to control protrusive activity through a myosin-dependent focal adhesion reinforcement that occurs distally of the protruding cell edge^20,21^. However, recent findings showed that localized Ca^2+^ influx promotes protrusive activity in macrophages on substrates that do not activate integrins^24^, indicating an alternative mechanism for Ca^2+^-driven lamellipodia formation. Here we demonstrate that TRPV4-mediated Ca^2+^ influx directly regulates the organization of branched actin network in protruding lamellipodia. We suggest that TRPV4-mediated Ca^2+^ signaling enhances the activity of formin-family actin regulators, facilitating the assembly of dense branched actin network. Indeed, formins Dia1 and Dia2 have previously been implicated in the assembly of branched actin network during lamellipodia protrusion^5,6^, and their inhibition has been shown to reduce Arp2/3 activation and suppress protrusive activity^17^, closely resembling the effect of TRPV4 suppression. Together, these data establish TRPV4 as a key local regulator of formin-mediated actin polymerization in lamellipodia.

Several studies have shown that formin activity is primarily regulated by small molecule GTPases, including RhoA^62,63^, suggesting a functional link between TRPV4 and RhoA signaling. A similar regulatory axis was recently described in neurons^64^, where the direct interaction between TRPV4 and RhoA leads to reciprocal inhibition^65^. However, our experimental results suggest a more complex, indirect regulation of RhoA by TRPV4 in protruding lamellipodia. Indeed, a quantitatively similar decrease in RhoA activity and actin density in lamellipodia of TRPV4-inhibited and TRPV4-depleted cells strongly indicates that the channel function, rather than direct interactions, controls RhoA activity. This conclusion is further supported by the effect of TRPV4 gain-of-function on RhoA activity in TRPV4-depleted cells. Immunofluorescence imaging reveals a uniform distribution of TRPV4 across the plasma membrane, with only marginal enrichment in lamellipodia. Despite this, TRPV4 overexpression selectively restores RhoA activity in lamellipodia, where Ca²⁺ influx was also re-established, suggesting the involvement of Ca²⁺-dependent signaling proteins in TRPV4-dependent RhoA activation.

The regulation of small GTPases by Ca^2+^ is a fundamental mechanism that governs the organization of actin cytoskeleton across various cell types^66–69^. However, molecular pathways that transduce Ca^2+^ signals to small GTPases remained elusive. Here, we identify two critical players that bridge Ca^2+^ signals in lamellipodia to RhoA activity - CaMKII and RhoGEF TEM4. Our findings demonstrate that Ca^2+^-dependent phosphorylation of TEM4 by CaMKII is essential for transducing TRPV4-mediated Ca^2+^influx to the activity of RhoA/formin actin polymerization machinery. Suppression of TRPV4 markedly reduces CaMKII activity and TEM4 phosphorylation, which correlates with decreased RhoA activity and less dense actin network. Similarly, CaMKII suppression produces the same phenotype. The essential role of a local pool of active CaMKII in the control of lamellipodial actin organization is further evident from the suppression of protrusions upon optogenetic inhibition of CaMKII. Notably, the expression of constitutively active CaMKII mutant or phospho-mimetic TEM4 variant fully rescues lamellipodial actin density in TRPV4 inhibited cells, suggesting that CaMKII-mediated TEM4 phosphorylation is sufficient to activate formin-mediated actin polymerization in lamellipodia. Interestingly, TEM4 is known to be autoinhibited via intramolecular interactions between its N- and C-termini^93^, and its phosphorylation may serve as a key regulatory mechanism to relieve this inhibition.

Our phosphoproteomic analysis also suggests an alternative pathway downstream of CaMKII that may contribute to lamellipodia actin regulation via LIM kinase (LIMK). We found that suppression of TRPV4 or CaMKII reduces phosphorylation of LIMK, a key regulator of actin severing protein cofilin. Therefore, activation of TRPV4/CaMKII signaling module may activate LIMK, which phosphorylates cofilin and inhibits its activity. Since phosphoregulation of cofilin is known to modulate lamellipodial actin^70,71^, this mechanism may work in concert with formins to regulate lamellipodia protrusion. Therefore, our data elucidate the first complete molecular pathway by which Ca^2+^ influx regulates small GTPase activity within a specific cellular domain – lamellipodia - and demonstrate the critical role of this regulatory mechanism in the control of actin organization.

More broadly, our study provides evidence that cells locally integrate Ca^2+^- dependent signaling and Rho family GTPases to fine-tune the organization of actin cytoskeleton. While we focused on understanding how the crosstalk between Ca^2+^ and GTPase signaling regulates F-actin organization in the context of cell migration, similar mechanisms may govern the assembly of actin cytoskeleton in other cellular processes, including membrane trafficking^72^ and cell-cell attachment^73^. Furthermore, this study identifies potential therapeutic targets - TRPV4, CaMKII, and TEM4 - for modulating cell migration in disease progression.

## LIMITATION OF THE STUDY

Our study reveals that TRPV4 is predominantly active in lamellipodia where it regulates CaMKII/TEM4/RhoA signaling module that converge on formin-mediated actin polymerization. But how does TRPV4 activity and the resulting Ca^2+^ influx are constrained in lamellipodia creating a local signaling hub? Several modes of TRPV4 regulation have been proposed, including activation by temperature changes^74^, mechanical stretching^75^, and membrane tension^76^. While temperature-driven activation is unlikely to control TRPV4 activity at the subcellular scale, gradients in membrane tension and cytoskeleton-generated frictional forces present plausible mechanisms for TRPV4 activation in lamellipodia.

Previous studies have shown that the pushing force generated by actin polymerization at the cell front builds up local plasma membrane tension^77,78^, providing the mechanical basis for the activation of mechano-gated ion channels. However, the rapid propagation of membrane tension along the plasma membrane, as reported previously^79^, would likely preclude localized activation of TRPV4 in protruding lamellipodia. Moreover, if membrane tension were the sole trigger, other mechanosensitive channels, such as TRPC1/6, TRPM7, Piezo^34^, would also be activated in lamellipodia. Instead, our data demonstrate that TRPV4 is the primary stretch-activated channel that governs the influx of Ca^2+^ in lamellipodia, suggesting a more specific regulatory mechanism of TRPV4 activation.

Based on the structural studies, we speculate that the specificity of TRPV4 activation in lamellipodia may stem from a unique actin-binding motif in its cytosolic C-terminus^80,81^. The motif may tether TRPV4 at the plasma membrane to the underlying actin network undergoing retrograde flow, thereby exerting a drag force on the channel molecules. In lamellipodia, where actin retrograde flow is rapid^82^, the resulting drag force may be sufficient to activate TRPV4, whereas the slower flow in the lamella may be insufficient, creating a sharp gradient of Ca^2+^ influx at cellular protrusions. Such spatially restricted TRPV4 activation, in turn, promotes formin-mediated polymerization and lamellipodia protrusion, establishing a positive feedback loop between formin activity and protrusion dynamics. Notably, such mechanochemical loop has been recently proposed^43^, but its molecular identity remains unclear. Our findings suggest that Ca^2+^ signaling may serve as the missing link in the mechanochemical regulation of actin assembly in the protruding cell edge, thereby facilitating cell migration.

## METHODS

### Cell Culture and Reagents

Mouse embryo fibroblasts (MEFs) were obtained from Dr. Beckerle (University of Utah). U-2 OS human osteosarcoma cells (U2OS) were obtained from Dr. Waterman (NIH). MDA-MB-231 and HeLa cells were a gift from Dr. Chen (Johns Hopkins University). MEFs and MDA-MB-231 cells were cultured in DMEM supplemented with 10% FBS, 2 mM L-glutamine, 100 U/mL penicillin/streptomycin. HeLa cells were cultured in DMEM supplemented with 10% FBS and 100 U/mL penicillin/streptomycin. All cells were maintained at 37°C in a humidified 5% CO_2_ incubator. All cell culture reagents were purchased from Wisent Bioproducts unless otherwise stated. The following chemicals and pharmacological inhibitors were used: DMSO (Fisher BioReagents, Cat: BP231100), BAPTA-AM (Cayman, Cat: 15551), hapsigargin (Cayman, Cat: 10522), EGTA (Bio Basic, Cat: ED0077), NP-EGTA-AM (Invitrogen, Cat: N6802), ionomycin (Millipore Sigma, Cat: I3909), GdCl_3_ (Millipore Sigma, Cat: 203289), HC-067047 (Millipore Sigma, Cat: 616521), C3 transferase (Cytoskeleton, Cat: CT04), ML-7 (Cayman, Cat: 11801), blebbistatin (Millipore Sigma, Cat: B0560), KN-93 (Cayman, Cat: 13319), SMIFH2 (Cayman, Cat: 31517).

### DNA Constructs

All of the plasmids generated for this study were created using restriction enzymes cloning. Primers were purchased from Integrated DNA Technologies (IDT), and the sequence of interest was confirmed by Sanger sequencing (TCAG DNA Sequencing Facility) and whole-plasmid sequencing (Plasmidsaurus). All oligonucleotide primers are listed in Table S2.

The stoichiometric calcium and cell volume reporter consists of cDNA encoding mCherry, a Gly-Ser-Gly linker, a T2A ribosomal skipping sequence, and GCaMP6f. The GCaMP6s insert and mCherry vector were PCR-amplified from GCaMP6s-CAAX (Addgene #52228) and mCherry-CAAX (Addgene #55045) using Phusion polymerase (NEB, Cat: M0530), BamHI/EcoRI-digested, and ligated with T4 ligase (NEB, Cat: MS0202). GCaMP6s in the assembled construct was subsequently modified to GCaMP6f by mutating A317 to E317^84^.

To generate a fluorescently labelled siRNA resistant TRPV4, cDNA encoding human TRPV4 (hTRPV4) from hTRPV4-TurboGFP (Origene #RG220160) was PCR-amplified by Phusion polymerase and XhoI/BamHI-digested. The digested PCR amplicon was ligated to a XhoI/BamHI-digested mEGFP vector, mEGFP-C1 (Addgene #54759), with T4 ligase.

To generate a far-red F-actin probe (F-tractin-emiRFP670), the F-tractin destination vector was created by PCR-amplifying the F-tractin-eGFP (Addgene #58473) while excluding the eGFP coding region using Phusion polymerase. The emiRFP670 insert from pemiRFP670-N1(Addgene #136556) was PCR-amplified using Phusion polymerase. The vector and insert were EcoRI/SbfI-digested and ligated with T4 ligase.

To create an overexpression construct of photoactivable CaMKII inhibitor (paAIP2) for MEFs, the histone H2B-eBFP plasmid (Addgene # 55243), which was used as the destination vector, and the cDNA region encoding a polycistronic P2A peptide and paAIP2 from CaMKIIP-mEGFP-P2A-paAIP2/pAAV (Addgene #91718) were PCR-amplified using Phusion polymerase. The vector and insert were NotI-digested and ligated with T4 ligase, resulting in the expression construct, H2B-eBFP2-P2A-paAIP2. A colony PCR screen was conducted to identify colonies that contain the plasmid with the correct insert orientation using Red-Taq polymerase (Wisent, Cat: 801-200).

To create an expression construct of FLAG tagged TEM4 with a cytoplasmic fluorescent marker as an indicator of expression, emiRFP670-T2A-3×FLAG::hTEM4 was generated in 2 sequential steps. First, emiRFP670 from F-tractin-emiRFP670 was PCR-amplified and HindIII/SbfI-digested. The digested PCR amplicon was ligated to a custom-made HindIII/SbfI -digested vector encoding hTEM4 (Genescript). Then, emiRFP670-hTEM4 was PCR-amplified with primers encoding T2A and 3×FLAG sequences. The PCR amplicon was SbfI-digested and ligated with T4 ligase, resulting in the final construct, emiRFP670-T2A-3×FLAG::hTEM4.

All the site-directed mutagenesis was performed based on a modified protocol^85^. Briefly, the mutagenesis primers were designed with 20 bps complementary sequences flanking the mutation site in both forward and reverse directions. Touchdown single-primer PCRs were performed with either the forward or reverse primer using Q5 polymerase (NEB, Cat: M0491) in the KOD polymerase buffer (Millipore Sigma, Cat: 71975). The forward and reverse PCR amplicons were subsequently combined, annealed, DpnI-digested, and transformed into competent cells (NEB, Cat: C2987).

Colonies were selected and screened for the presence of the desired mutation sites by Sanger sequencing (TCAG DNA Sequencing Facility) and subsequently whole-plasmid sequencing (Plasmidsaurus).

### Cell Transfection with pDNA Expression Constructs and siRNAs

For experiments that require transient expression of recombinant protein(s) or siRNA-mediated protein depletion, cells were transfected with 1 μg of a desired pDNA expression construct(s) and/or 200 nM siRNAs in 100 μL of a house made transfection solution (5 mM KCl, 15 mM MgCl_2_, 120 mM Na_2_HPO_4_/NaH_2_PO_4_ (pH 7.2), 50 mM mannitol). Cells were transfected using Nucleofector 2b (Lonza Bioscience) with program T-020. Transfected cells were plated on glass-bottom imaging dishes (Cellvis, Cat: D35-20-1.5-N) or coverslips that were coated with 2.5 μg/mL human fibronectin (EMD Millipore, Cat: FC010) in PBS for 1 hr at 37°C.

### Immunofluorescence and Phalloidin Staining

Cells seeded on fibronectin-coated coverslips (Corning) were fixed in 4% EM-grade paraformaldehyde (Electron Microscopy Sciences, Cat: 15700) diluted in cytoskeleton buffer (10 mM MES (pH 6.1), 138 mM KCl, 3 mM MgCl_2_, 2 mM EGTA, 0.32 M sucrose) for 30 min at room temperature and washed 3 times with PBS. After fixation, cells were permeabilized with 0.1% (v/v) Triton X-100 (BioShop, Cat: TRX777) in PBS for 10 min, washed with PBS and blocked in 2% (w/v) BSA (BioShop, Cat: ALB001) dissolved in PBS. Cells on coverslips were incubated overnight at 4°C with primary antibodies (1:100) diluted in the blocking solution. The primary antibodies used for immunofluorescence in this study were: anti-ArpC2 (Millipore Sigma, Cat: 07-227), anti-cortactin (Cell Signaling, Cat: 3503S), anti-pS19MLC2 (Cell Signaling, Cat: 3674S), anti-eGFP (Cell Signaling, Cat: 2555S), anti-pCaMKII (Abcam, Cat: ab32678). Following the incubation with the primary antibodies, the coverslips were washed with PBS supplemented with 0.1% Tween-20 (BioShop, Cat: TWN510) and incubated for 1 hr with fluorophore-conjugated secondary antibodies (1:200, Jackson ImmunoResearch Laboratories) and Alexa Fluor488-conjugated phalloidin (ThermoFisher, Cat: A12379) in the blocking solution. The stained samples were washed with PBS containing 0.1% Tween-20, mounted on microscope slides in an aqueous mounting media (Fluoromount, Sigma-Aldrich, Cat: F4680) and sealed with nail polish.

### Microscopy and Live Cell Imaging

All live-cell imaging experiments were conducted at 37°C and 85% relative humidity using a stage-top incubator and an objective heater (Tokai Hit). Cells were plated on fibronectin-coated coverslips and imaged in a DMEM imaging media, composed of phenol red-free DMEM, 25 mM HEPES, 10% FBS, 100 U/mL penicillin/streptomycin, 10 mM DL-lactate (Millipore Sigma, Cat: 71720), and OxyFluor (Oxyrase, Cat: OF-0005), unless otherwise stated.

Images of live and immunolabeled cells were acquired on an Eclipse Ti2-E inverted microscope (Nikon Instruments) equipped with a phase contrast 10×/0.3 NA Plan Fluor DLL, 20×/0.75 NA Super Fluor, 100×/1.45 NA CFI Plan Apochromat Lambda D, and 100×/1.49 NA CFI Apo TIRF objectives, a dynamic focusing system to correct for focus drift (PFS, Nikon Instruments), a CREST X-Light V2 LFOV25 (CrestOptics), a 7-color Celesta light engine (Lumencor) and a Prime95b sCmos camera (Photometrics).

For the light stimulation imaging experiments, images of cells were acquired on an Eclipse Ti2-E inverted microscope (Nikon Instruments) equipped with a 60×/1.49 NA CFI Apo TIRF objective, a dynamic focusing system to correct for focus drift (PFS; Nikon Instruments), a CSU-X1 confocal scanner (Yokogawa Corporation) with 4-color iChromeMLE laser combiner (Toptica Photonics), a CoolSnap Myo CCD camera (Roper Scientific), and a dual-galvanometer laser scanner (Bruker Corporation) with a 100mW 405/488 nm LunF solid state laser illumination unit (Nikon Instruments).

Images of cells cultured in three-dimensional collagen gel were acquired by an Eclipse FN1 upright microscope (Nikon Instruments) equipped with an A1R-MP confocal scanner, a femtosecond infrared laser (Chameleon Vision II, Coherent) and 25×/1.1 NA CFI75 Apo water immersion objective. Second harmonic generation signal emitted by collagen fibers was collected with a non-descanned GaAsP epi-detector after it was isolated from the laser fundamental and any fluorescence by a 474 nm bandpass filter (28 nm full width half-maximum; Semrock Inc, Cat: FF-01-474/23 nm BrightLine).

### Reverse Transcription PCR (RT-PCR)

Total RNA was isolated from MEFs with PureLink RNA Mini kit (Invitrogen, Cat: 12183018A) and treated with DNase I (Invitrogen, Cat: 18047019) to remove DNA contamination. cDNA was synthesized using random hexamer primers with SuperScript II Reverse Transcriptase (Invitrogen, Cat: 18064-022). To amplify the cDNA, PCR was performed with DreamTaq PCR Master Mix (Thermo Scientific, Cat: K1081). PCR primers were designed on Primer3. The primer sequences are listed in Supplementary Table 1.

### Western Blotting

Cell lysates were prepared from the cultured cells after removing the growth media and rinsing with ice-cold PBS. Cells were lysed directly by scraping in RIPA buffer (150 mM NaCl, 1% (v/v) Nonidet P-40, 0.5% (w/v) sodium deoxycholate, 0.1% (w/v) SDS, 50 mM Tris-HCl (pH 7.4)) supplemented with a protease inhibitor cocktail (Roche, Cat: 11697498001) and a phosphatase inhibitor cocktail (Roche, Cat: 4906845001). The lysates were sheared by running through a 25G needle 10 times and then cleared by centrifugation (17,900 ×*g*, 25 min, 4°C). Protein concentration in the supernatants was measured by the BCA protein assay (ThermoFisher, Cat: 23225). Aliquots of cell lysates containing 40 µg of proteins were mixed with an equal volume of 2× Laemmli sample buffer and separated by SDS-PAGE on 8% or 12% gels in Tris-glycine running buffer (Wisent, Cat: 811-560-FL) at 30 mA and then electro-transferred to PVDF membranes (Millipore Sigma, Cat: ISEQ10100) at 35 V overnight or at 90 V for 60 min. To facilitate the overnight transfer of TEM4, CAPS buffer (0.5 M 3-(Cyclohexylamino)-1-propanesulfonic acid, pH 11, 10% methanol) was used as the transfer buffer. Following transfer, the membranes were blocked for 1 hr. For fluorescence detection, TBS blocking buffer (Licor, Cat: 927-60001) was used. For chemiluminescence detection, 5% (w/v) BSA (Wisent) in TBST (Wisent, Cat: 811-030-FL, 1× buffer supplemented with 0.1% (v/v) Tween-20) was used. After blocking, the membranes were incubated with primary antibodies, followed by a wash and incubation with secondary antibodies conjugated with an IRDye or HRP for fluorescent and chemiluminescent detection, respectively.

Fluorescently labelled membranes were washed with TBST and imaged with the Odyssey® Fc imaging system (LI-COR). HRP-conjugated secondary antibodies were visualized using enhanced chemiluminescent horseradish peroxidase (HRP) substrate (ThermoFisher, Cat: 34580) and imaged with the ChemiDoc MP system (BioRad). The antibodies and dilutions used for western blotting in this study were: anti-TRPV4 (1:1000, Abcam, Cat: ab39260), anti-vimentin (1:2000, Cell Signaling, Cat: 5741S), anti-RhoA (1:500, Cytoskeleton, Cat: ARH05), anti-Rac1 (1:500, Cytoskeleton, Cat: ARC03), anti-Cdc42 (1:250, Cytoskeleton, Cat: ACD03B), anti-pCaMKII (1:1000, Abcam, Cat: ab32678), anti-vinculin (1:1000, MilliporeSigma, Cat: V9131), anti-VASP (1:1000, Cell Signaling, Cat: 3132T), anti-TEM4 (1:500, Novus Biologicals, Cat: NBP1-77314).

For chemiluminescence detection, HRP-conjugated donkey anti-mouse IgG or anti-rabbit IgG purchased from Jackson Immuno Research were used as secondary antibodies. The HRP-conjugated antibodies were diluted 1:10,000 in TBS supplemented with 0.1% (v/v) Tween-20 and 2% BSA. For fluorescent antibody detection, IRDye (680 nm or 800 nm)- conjugated anti-mouse IgG or anti-rabbit IgG were used at a dilution 1:10,000 in TBS blocking buffer (LI-COR). Visualization and quantification of western blots was performed using ImageStudio (LI-COR).

### Platinum Replica Electron Microscopy (PREM) and EM Image Analysis

Sample processing for PREM was performed as described previously^86,87^. In brief, cells were seeded on glass coverslips coated with 2.5 μg/mL fibronectin. After 1 hr of drug treatment in culture, cells were extracted for 4 min with 1% Triton X-100 in PEM buffer (100 mM PIPES, pH 6.9; 1 mM EGTA; 1 mM MgCl_2_) supplemented with 2 μM phalloidin and 2 μM taxol and fixed with 2% glutaraldehyde in 0.1 M Na-cacodylate buffer (pH 7.3) for 20 min. The samples were post-fixed by sequential treatment with aqueous 0.1% tannic acid and 0.2% uranyl acetate, then critical-point dried, coated with platinum and carbon, transferred onto EM grids and examined using JEM 1011 transmission electron microscope (JEOL USA, Peabody, MA) operated at 100 kV. Images were acquired by an ORIUS 832.10 W CCD camera driven by Gatan Digital Micrograph 1.8.4 software (Gatan, Warrendale, PA) and presented in inverted contrast.

The EM images were processed and analyzed using a custom-built Python script (skeleton_EM.py) and Fiji/ImageJ. Briefly, a Gaussian blur filter was applied to the EM images to reduce noise, followed by a top-hat transformation to extract the linear features of the images. The preprocessed images were binarized using the K-means clustering algorithm and the fraction of background pixels, representing areas devoid of F-actin, was calculated and reported as the porosity of the actin network. Pore size quantification was performed using inverted binary maps of F-actin and Analyze Particles tool in Fiji/ImageJ.

### Cell Spreading Assay and Cell Spreading Speed Measurement

The cell spreading assay was performed as previously described^88^. Briefly, cells were harvested using 0.05% trypsin-EDTA, resuspended in cell culture medium supplemented with 10% FBS, and allowed to recover in suspension for 45 min at 37°C. Approximately 50,000 cells/mL were then plated onto a fibronectin-coated 22-by-22 mm coverslip mounted in a Chamlide magnetic chamber (Quorum Technologies), and phase contrast images of the spreading cells were acquired at 1 sec interval for 10 min using 10×/0.3 NA Plan Fluor DLL objective. To quantify cell spreading, we generated kymographs from the spreading cell edges using the KymoResliceWide plugin in Fiji/ImageJ. The slope of fast spreading phase was reported as the average spreading speed. To compare data from different experimental treatments and biological replicas, the values from each experiment were normalized to the mean cell spreading speed of the corresponding control.

### Lamellipodia Protrusion Speed Measurement

MEFs were plated on fibronectin-coated coverslips 12 hr prior to the experiment. To visualize cell edge dynamics, time-lapse DIC images were acquired at 3 sec interval for 20 min using 100×/1.49 NA CFI Apo TIRF objective. To quantify protrusion velocity, a three-pixel-wide line was placed normal to the free cell edge, and a kymograph was generated using the KymoResliceWide plugin in Fiji/ImageJ. Lamellipodia protrusion velocity was calculated from the kymographs, and the mean protrusion speed for each cell was determined by averaging the slopes of five protrusion events.

### Measurements of Lamellipodial F-Actin, Arpc2, Cortactin, pCamkII Intensities, and Arpc2-enriched Cell Perimeter

MEFs co-stained with phalloidin and anti-cortactin, anti-pCaMKII, or anti-ArpC2, were imaged using a spinning disk confocal microscope. To account for local changes in image intensity due to membrane ruffles near cell edges, 1.2 µm confocal z-stacks were captured. The acquired stacks were processed by maximum intensity projection and background subtraction using a custom Python script (background_subtraction_nd2_images.py). To quantify F-actin and lamellipodia marker intensities, ROIs were manually outlined around lamellipodia in the processed images using Fiji/ImageJ. The mean grayscale value within the ROI was reported as the F-actin, ArpC2, or cortactin intensity.

The percent of ArpC2-enriched perimeter was determined by measuring the length of ArpC2-enriched (designated as intensity approximately greater than 1.5 standard deviations above the mean intensity for the whole cell) cell perimeter and then dividing it by the total cell perimeter.

### Calcium Imaging

For simultaneous visualization of intracellular Ca^2+^ and F-actin, MEFs were co-transfected with mCherry-T2A-GCaMP6f and F-tractin-emiRFP670 expression constructs and cultured for 24 hr. Transfected cells were treated with DMSO (vehicle) or the desired pharmacological inhibitor 1 hr before imaging. For spatiotemporal analysis of intracellular Ca^2+^ in spreading fibroblasts, transfected cells were plated on coverslips coated with 2.5 µg/mL fibronectin, and multi-color confocal movies were acquired at

1 sec interval for 10 min using a 100×/1.45 NA CFI Plan Apochromat objective. To analyze Ca^2+^ dynamics in migrating cells, transfected fibroblasts were cultured on fibronectin-coated coverslips for 12 hr, and multi-color confocal movies were acquired at 10 sec intervals for 20 min. To observe the acute effects of changes in the intracellular Ca^2+^, transfected cells were plated on glass-bottom dish coated with 2.5 µg/mL fibronectin and imaged every 10 sec for 5 min in the presence of DMSO. An equal volume of imaging media supplemented with 20 μM HC-067047 or 2 μM ionomycin was then added and the imaging continued for an additional 15 min.

### RhoA/Rac1/Cdc42 Activity Assay

A Rho GTPase activation biochemical assay (Cytoskeleton, Cat: BK030) was used to measure RhoA, Rac1, and Cdc42 activity, following the manufacturer’s recommendations. MEFs were cultured to 80% confluency and then treated for 1 hr with either 0.1% DMSO (vehicle), 10 μM HC-067047, or 10 μM KN-93. The cells were then washed with ice-cold PBS and lysed in 250 μL of sample lysis buffer. Samples containing 350 μg of protein were incubated with rhotekin-RBD beads and 800 μg of protein with PAK-PBD beads for 3 hr, washed, and analyzed by Western blotting.

### RhoGTPase FRET Biosensors Imaging

To assess the subcellular activity of GTPases (RhoA, Rac1 and Cdc42), MEFs were transfected with pTriExRhoA2G (Addgene #40176), pTriEx4-Rac1-2G (Addgene #66110), or pTriEx4-Cdc42-2G (Addgene #68814), respectively. Transfected cells were plated into glass-bottom dishes coated with 2.5 µg/mL fibronectin and cultured overnight. 1 hr prior to imaging, the cell culture medium was replaced with live cell imaging solution (Invitrogen, Cat: A59688DJ), supplemented with 2% FBS, 100 U/mL penicillin/streptomycin, 10 mM HEPES, 10 mM DL-lactate, and OxyFluor. For experimental conditions requiring TRPV4 and CaMKII inhibition, cells were treated for 1 hr with 10 μM HC-067047 and 10 μM KN-93, respectively. Time-series of widefield images of the donor, acceptor and FRET channels were acquired using a 60×/1.2 NA Plan Apo VC water immersion objective every 3 min for 20 min.

### Ratiometric Image Analysis

The ratiometric image analysis was performed according to a published pipeline^89,90^ using custom-built Python scripts (get_ratiometric_map.py for GCaMP6f images and FRET_get_ratiometric_map.py for images of the RhoGTPase biosensors).

Briefly, multi-channel images were pre-processed by applying shading correction with a cell-free reference image, followed by alignment using the optical flow registration algorithm. The intensity of images was adjusted by subtracting the median intensity of the background pixels, determined by the Otsu method. For FRET image analysis, a bleed-through correction was applied. The images were denoised by applying a gaussian filter (σ = 0.5) and ratiometric images were generated by dividing the reporter (GCaMP or FRET) channel by the reference (mCherry or mTFP1) channel. Spurious pixels outside the cell were removed by applying logical AND to the ratiometric image and a binary cell mask, generated from the corresponding F-tractin or mVenus channel.

For the FRET image analysis, the following equation was used to compensate for the light bleed-through:

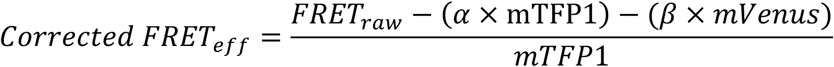

where FRET_raw_ and Corrected FRET_eff_ are the total and corrected FRET efficiency, respectively, α is the bleed-through coefficient of mTFP1 into the FRET channel upon mTFP1 (440 nm) excitation, β is the bleed-through coefficient of mVenus into FRET channel upon mVenus (514 nm) excitation, and mTFP1 is the total mTFP1 intensity as measured. MEFs transiently expressing either mTFP1 or mVenus were used to determine α and β. Consistent with previous reports^89,90^, the measured values of the α and β coefficients were 0.3072 and 0.05228, respectively.

### Correlation analysis of cell edge displacement and Ca^2+^ level

The cross-correlation analysis was performed as described previously^91^. To obtain the spatial and temporal information of cell edge displacement and Ca^2+^ activity, kymographs based on the ratiometric Ca^2+^ images were created using the KymoResliceWide plugin in Fiji/ImageJ (averaging 3 pixels in width). Then, each kymograph was imported into and analyzed by a custom Python script (quantify_cell_edge_velocity_and_calcium.py). To compute the cross-correlation of cell edge displacement and Ca^2+^ level, the temporal information of cell edge displacement and lamellipodial Ca^2+^ level were required. A cell edge displacement-time plot was extracted from the kymograph by tracking the cell edge position over time. A Ca^2+^-time plot was extracted by measuring the average intensity of the 4 pixels (∼1.5 µm) closest the cell edge. These two plots were Gaussian-filtered (σ = 1) before the cross-correlation analysis. The cross-correlation coefficient was computed by the np.correlate function and the delay times by the scipy.signal.correlation_lags function. A spline fitting was applied to smoothen the curve in the cross-correlation graph.

### Photo-release of Caged Ca^2+^

MEFs were transfected with mCherry-T2A-GCaMP6f, plated in glass-bottom dishes coated with 2.5 µg/mL fibronectin, and cultured overnight. The cells were treated with 10 µM NP-EGTA-AM dissolved in DMSO with 0.1% Pluronic for 2 hr, after which the cell culture medium was replaced with the imaging media supplemented with 1 µM myosin light chain kinase inhibitor ML-7. The inhibition of myosin light chain kinase prevented cell retraction caused by Ca^2+^-dependent myosin activation in the lamella^21^.

Small Ca^2+^ pulses were generated by illuminating a 3×3 µm region of the cytoplasm adjacent to the stalling cell edge with a low-intensity 405 nm laser for 3 sec. Time-lapse DIC images were captured every 3 sec for 30 sec before Ca^2+^ uncaging and for 2 min after the uncaging. Time-lapse GCaMP6f and F-tractin-emiRFP670 images were captured every 15 sec for 30 sec before Ca^2+^ uncaging and for 2 min after the uncaging.

### CaMKII Photoinactivation

For local suppression of CaMKII activity, we used an optogenetic inhibitor paAIP2^47,50^. MEFs were transfected with the H2B-eBFP2-P2A-paAIP2 expression construct, plated in glass-bottom dishes coated with 2.5 μg/mL fibronectin, and cultured overnight. 1 hr prior to the experiment, the cell culture media was replaced with the imaging media, samples were visually inspected and H2B-eBFP2 positive cells were selected for the experiments. The optogenetic inhibitor of CaMKII was activated by illuminating a 3×3 µm region of the cytoplasm directly adjacent to the protruding cell edge with a low-intensity 405 nm laser for 3 sec. Timelapse DIC images were captured every 3 sec for 30 sec before paAIP2 activation and for 2 min after the activation.

### Isolation of Lamellipodia and Cell Body Fractionation

The lamellipodia and cell body fractionation was performed as described^54^. Briefly, MEFs were plated on a 3-µm porous polycarbonate membrane (Millipore Sigma, Cat: CLS3464) pre-coated with fibronectin. After cell attachment, the cells were starved overnight in the media supplemented with 1% FBS. The starved cells were subjected to a serum gradient (1% in the inner compartment and 10% in the outer compartment) overnight followed by methanol fixation. The lamellipodia fraction was first harvested manually with a cotton swab. The cotton swab was subsequently placed in RIPA buffer to solubilize the proteins. Then, the cell body fraction was extracted by placing the remaining membrane in RIPA buffer. Protein concentration in all samples was measured by the BCA assay. To validate the fractionation of lamellipodia and cell body, we routinely compare the abundance of VASP, as a lamellipodia marker, in lamellipodia and cell body fractions using Western blot.

### Phospho-peptide enrichment and mass spectrometry

Sample preparation and analysis were performed as previously described^92,93^. Lamellipodia (LP) or cell body (CB) fractions (400 μg) were reduced (15 mM DTT, 56°C, 45 min) and alkylated (50 mM iodoacetamide, room temperature, 45 min). Proteins were precipitated (4× volume acetone, -20°C, 16 hr), collected by centrifugation (16,000 ×g 4°C, 20 min) and resuspended in 0.1% (w/v) Rapigest (Waters) in 100 mM ammonium bicarbonate. Proteins were digested with trypsin at a 1:50 (w/v) ratio (37°C, 16 hr with shaking at 250 rpm) and then acidified with 1% (v/v) trifluoracetic acid (TFA; 37°C ,45 min). Prior to phospho-peptide enrichment samples were desalted using Oasis HLB cartridges (Waters) according to manufacturer’s instructions, eluting in binding solution: 80% (v/v) acetonitrile (ACN), 5% (v/v) TFA, 1 M glycolic acid. For proteomic analysis 5% of the eluted sample was retained and stored at -20°C.

Phospho-peptide enrichment was performed using magnetic microspheres (Ti-IMAC and TiO2; ReSyn Bioscience) and a KingFisher Flex (Thermo Scientific) magnetic particle-processing robot^94^. For each sample 800 μg microspheres (25% : 75% Ti-IMAC : TiO_2_ ratio) were prepared according to manufacturer’s instructions before incubation with tryptic digests for 20 min under agitation. Following washes in 80% (v/v) ACN / 1% (v/v) TFA and 10% (v/v) ACN / 0.2% (v/v) TFA, peptides were eluted in 1% (w/v) ammonium bicarbonate for 10 min and acidified to 1% (v/v) with TFA. Three rounds of phospho-peptide enrichment were performed and elutions pooled for each sample. Phospho-peptides were desalted using Poros R3 beads (Applied Biosystems, UK), collected by elution in 50% (v/v) ACN / 0.1% (v/v) formic acid, dried under vacuum and stored dried at 4°C.

For mass spectrometry peptides were resuspended in 5% (v/v) ACN / 1% (v/v) formic acid and analysed by liquid chromatography (LC) tandem mass spectrometry (LC-MS/MS) using an UltiMate 3000 Rapid Separation LC (RSLC, Dionex Corporation) coupled to a Q Exactive HF (Thermo Fisher Scientific) mass spectrometer. Peptides were separated using a 75 mm × 250 μM i.d. 1.7 μM CSH C18, analytical column (Waters) with the LC gradient from 95% buffer A (0.1% (v/v) FA in water) and 5% buffer B (0.1% (v/v) FA in acetonitrile) to 7% buffer B at 1 min, 18% buffer B at 58 min, 27% buffer B in 72 min and 60% buffer B at 74 min at 300 nL min^-1^. Peptides were selected for fragmentation automatically by data dependant analysis. For phosphoproteomic analyses, multistage activation was enabled to fragment product ions resulting from neutral loss of phosphoric acid.

Relative protein and phospho-peptide abundances were calculated based on ion intensity as implemented in Proteome Discoverer (v2.3; ThermoFisher Scientific). RAW files were searched using SEQUEST-HT against the SwissProt and TREMBL mouse databases (release-2018_01). Carbamidomethylation of cysteine was set as a fixed modification; serine, threonine and tyrosine phosphorylation and oxidation of methionine were allowed as variable modifications. Mass tolerances for precursor and fragment ions were 10 ppm Da and 0.02 Da, respectively. Abundance ratios were calculated between treatment groups, together with adjusted P-values using the Benjamini-Hochberg method, in Proteome Discoverer.

### Random migration assay and cell tracking

For automated unbiased cell tracking, MEFs were fluorescently labeled by expressing histone H2B tagged with mEmerald and sparsely seeded onto 2.5 µg/mL fibronectin-coated coverslips. The coverslips with cells were mounted in a Chamlide CMS magnetic chamber (Quorum Technologies) 24 hr after plating. Timelapse DIC and fluorescence images were acquired through a 20×/0.75 NA Super Fluor objective every 12 min for 20 hr.

The timelapse images of fluorescently labelled nuclei were analyzed using the Fiji/ImageJ plugin TrackMate. Nuclei were first identified using the LoG detector (estimated object diameter: 25 µm, quality threshold: 0.08). The LAP tracker algorithm (max distance: 50 µm, Gap closing max distance and max frame gap: 50 µm and 3 frames) was then applied to track nuclei movement in the cell movies. The resulting cell tracks were filtered based on duration and length, restricting the analysis to tracks that lasted at least 60 min and spanned a minimum of 18 µm. Tracks were manually terminated once the cell began to divide. Temporal and positional data for each track were exported as comma-separated values for further computational analysis and visualization using a custom-built Python script (trackmate_csv.py).

### Collagen matrix preparation, morphological and cell migration analysis in 3D microenvironment

Collagen matrices were prepared according to published protocols^95^. Briefly, 35-mm glass bottom dishes were first silanized by (3-aminopropyl) trimethoxysilane (APTMS, Millipore Sigma, Cat: 281778) and activated with glutaraldehyde (Electron Microscopy Sciences, Cat: 16220). Then, MEFs were mixed with neutralized collagen I (3 mg/mL) to achieve a cell density of 50 × 10^3^ cells/ml and 500 µL aliquots were added to the activated dishes. The cell/collagen mixture was allowed to polymerize at 21°C for 1 hr. After the collagen sponge formed, the dishes were transferred to a humidified incubator at 37°C with 5% CO₂ overnight, prior to imaging.

For the morphological analysis of cells embedded in collagen gels, MEFs were either pre-treated with 0.1% DMSO or 10 µM HC-067047 for 1 hr, or transfected with siRNAs (siCtrl or siTEM4) 72 hr prior to imaging. High magnification confocal z-stacks of individual cells were acquired by a CSU-X1 spinning disk confocal microscope (Yokogawa Corporation) paired with an oil immersion 60×/1.49 NA Apo TIRF objective. The average number of dendritic protrusion and total protrusion length were analyzed in NIS-Elements software (Nikon Instruments).

For the migration assay in 3D collagen gel, the nuclei of MEFs were labelled with 50 ng/mL Hoechst 33342 (Life Technologies, Cat: R37605) and cell migration was monitored over 12 hr. Confocal z-stacks were captured every 7 µm over a z-distance of 308 µm. To analyze the cell displacement, maximum intensity projected (MIP) images for each time point were first created and the cell movement was manually tracked from the MIP images in Fiji/ImageJ. Cells that migrated in the z direction were not included in the quantification. The 3D migration speed was calculated by dividing the cell displacement by the duration of the cell track.

### Ex vivo visualization of melanoblast migration

To label embryonic melanoblasts, a melanocyte-specific *Mitf-Cre* transgene was used to induce *tdTomato* expression from a *Rosa26-floxed stop-tdTomato* fluorescent reporter. E15.5 embryos were harvested, and the intact epidermis and dermis were isolated from the embryonic trunk skin. The skin explant was then sandwiched so that the epidermis was in contact with a gas permeable Lumox membrane in a 35 mm dish and the dermis was in contact with a 1% agarose pad made with complete phenol-free DMEM. Finally, the sandwich setup was overlaid with 1 mL complete phenol-free DMEM and mounted with a light weight for imaging. For the TRPV4 inhibitor treatment group, the skin explants were incubated with complete phenol-free DMEM with 10 µM HC-

067047 for 2 hr before imaging. Skin explants were imaged live using a 20× objective with a stage top incubator at 37°C on an Olympus Fluoview (FV1000) inverted confocal microscope.

Melanoblasts movies were analyzed using the Fiji/ImageJ ADAPT plugin, and single cells were tracked throughout the duration of the entire movie.

### Focal Adhesion Imaging and Image Analyses

To visualize focal adhesions and F-actin in MEFs, we fixed and co-stained cells with anti-vinculin (1:100; MilliporeSigma, Cat: V9131) and fluorescent phalloidin (Invitrogen, Cat: A12380). Fluorescence images of vinculin and F-actin were acquired through a CSU-X1 spinning disk confocal microscope (Yokogawa Corporation) paired with an oil immersion 60×/1.49 NA Apo TIRF objective.

The analysis pipeline of focal adhesion was performed as described previously^96^. Briefly, the area of individual focal adhesions was measured from the immunofluorescence images of cells stained with anti-vinculin antibody using the publicly available Focal Adhesion Analysis Server^97^. The classification of focal adhesions based on area was performed as follows: small vinculin-positive adhesions (area ranges from 0.05 to 0.20 µm^2^) were classified as nascent adhesions^8^. FAs with an area ranging from 0.20 to 1.75 µm^2^ and larger than 1.75 µm^2^ were classified as mature and fibrillar adhesions, respectively. The percentage of each type of adhesions was calculated for individual cells and then averaged for every experimental condition.

### Quantification and Statistical Analysis

Statistical analyses were performed with GraphPad Prism software. Details of statistical operations performed in figures are presented in their corresponding figure legend.

## ACKNOWLEDGMENTS

We thank Drs. Tony Harris and Rudolf Winklbauer (University of Toronto) for insightful comments on the manuscript. We thank Dr. Sabine Elowe and Chantal Garand (University of Laval) for help with the analysis of TEM4 expression, and Steven Chen (University of Toronto) for providing technical advice on molecular cloning. This work was supported by Natural Sciences and Engineering Research Council of Canada discovery grants RGPIN-2015-05114 and RGPIN-2020-05881 (S.V.P.), Canadian Institutes of Health Research project grant PJT-178272 (S.V.P.), the National Institutes of Health grant R01GM148459 (S.V.P.), Cancer Research UK Programme Grant (DRCRPG-100002 to M.J.H.), and University of Manchester-University of Toronto Joint Research Fund (S.V.P. and M.J.H.). The mass spectrometers used in this study were purchased with grants to M.J.H. from BBSRC, Wellcome Trust and the University of Manchester Strategic Fund and the work was conducted within the Wellcome Centre for Cell-Matrix Research (core award 226804/Z/22/Z). E.I. was support by Ontario Graduate Scholarship and Natural Sciences and Engineering Research Council of Canada (NSERC PGS-D).

## SUPPLEMENTAL FIGURES

**Figure S1.**
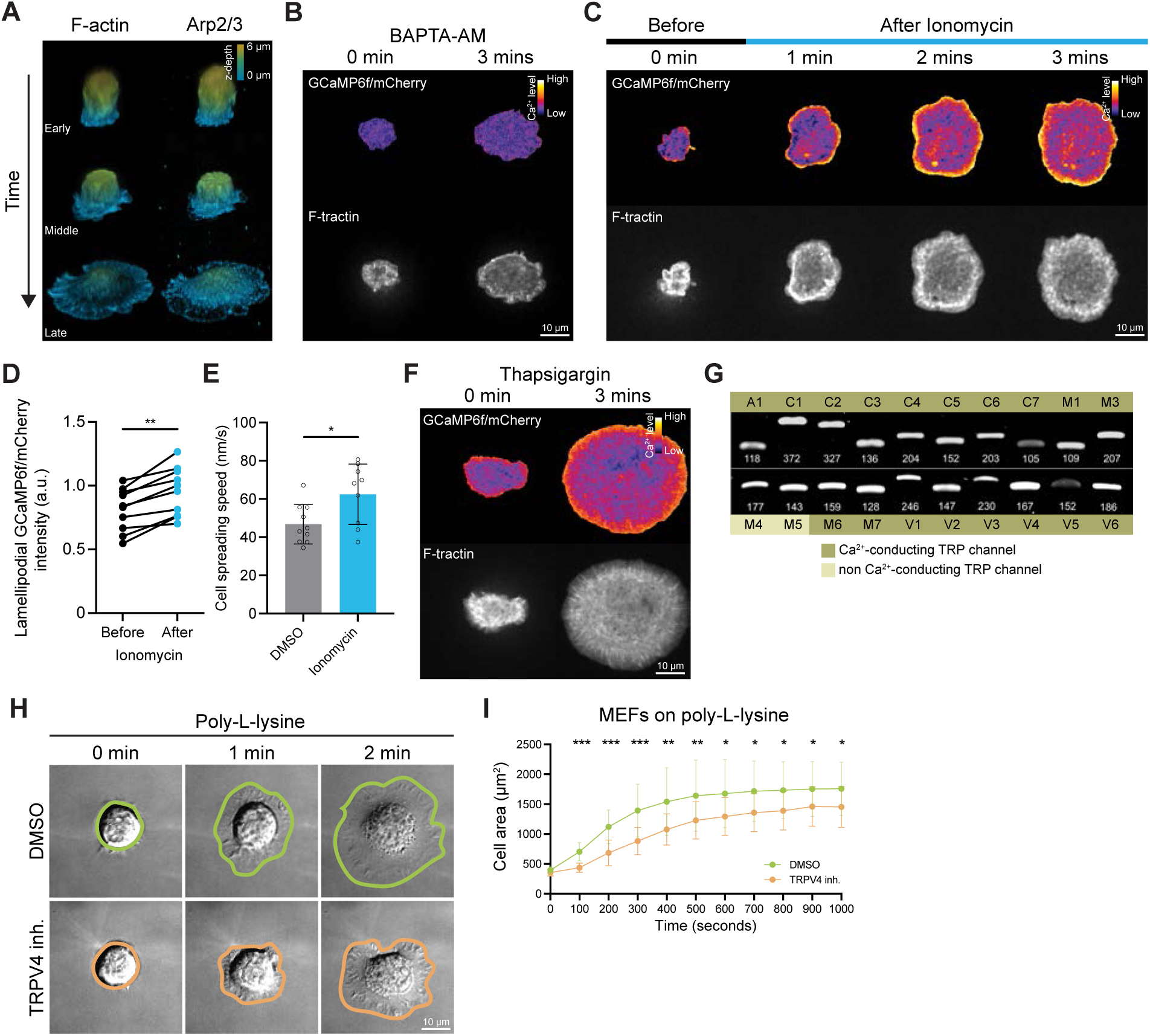
Perturbations in Ca^2+^ signaling, by Ca^2+^ drugs or TRPV4 inhibitor, affects cell spreading independently of integrin signaling. Related to Figure 1. (A) 3D projection of a spreading MEF in 3 stages (early, middle and late). (Left) Fluorescent phalloidin and (Right) anti-ArpC2 staining. The images are color-coded based on the z-depth. (B-F) Effects of Ca^2+^ ionophore, chelator and endoplasmic reticulum Ca^2+^ ATPases (SERCA pumps) inhibitor on cell spreading. Timelapse ratiometric GCaMP6f/mCherry and F-tractin-emiRFP670 images of a spreading cell treated with (B) 10 μM BAPTA-AM and (C) 1 μM ionomycin. Quantifications of (D) the lamellipodial GCaMP6f/mCherry intensity and (E) the average cell spreading speed before and after the addition of ionomycin. n ≥ 9 cells per condition. (F) Timelapse ratiometric GCaMP6f/mCherry and F-tractin-emiRFP670 images of a spreading cell treated with 1 μM thapsigargin. (G) Reverse transcription PCR (RT-PCR) analysis of TRP family channels expressed in MEFs. The TRP channels that are labelled in dark and light green are Ca^2+^-conducting and non Ca^2+^-conducting, respectively. (H-I) Effects of poly-L-lysine coated substrate on cell spreading. (H) DIC timelapse images of spreading MEFs treated with (top) DMSO and (bottom) TRPV4 inhibitor. The colored outline shows the cell boundary. (I) Quantifications of cell spread area over time in DMSO and TRPV4 inhibitor conditions. n = 20 cells per condition. Mann-Whitney test is used for comparisons between 2 treatments and one-way ANOVA followed by Tukey-Kramer post-hoc test for comparisons between 3 or more treatments. The data are presented as mean ± standard deviation (SD). * p < 0.05, ** p < 0.01, *** p < 0.005, **** p < 0.001.

**Figure S2.**
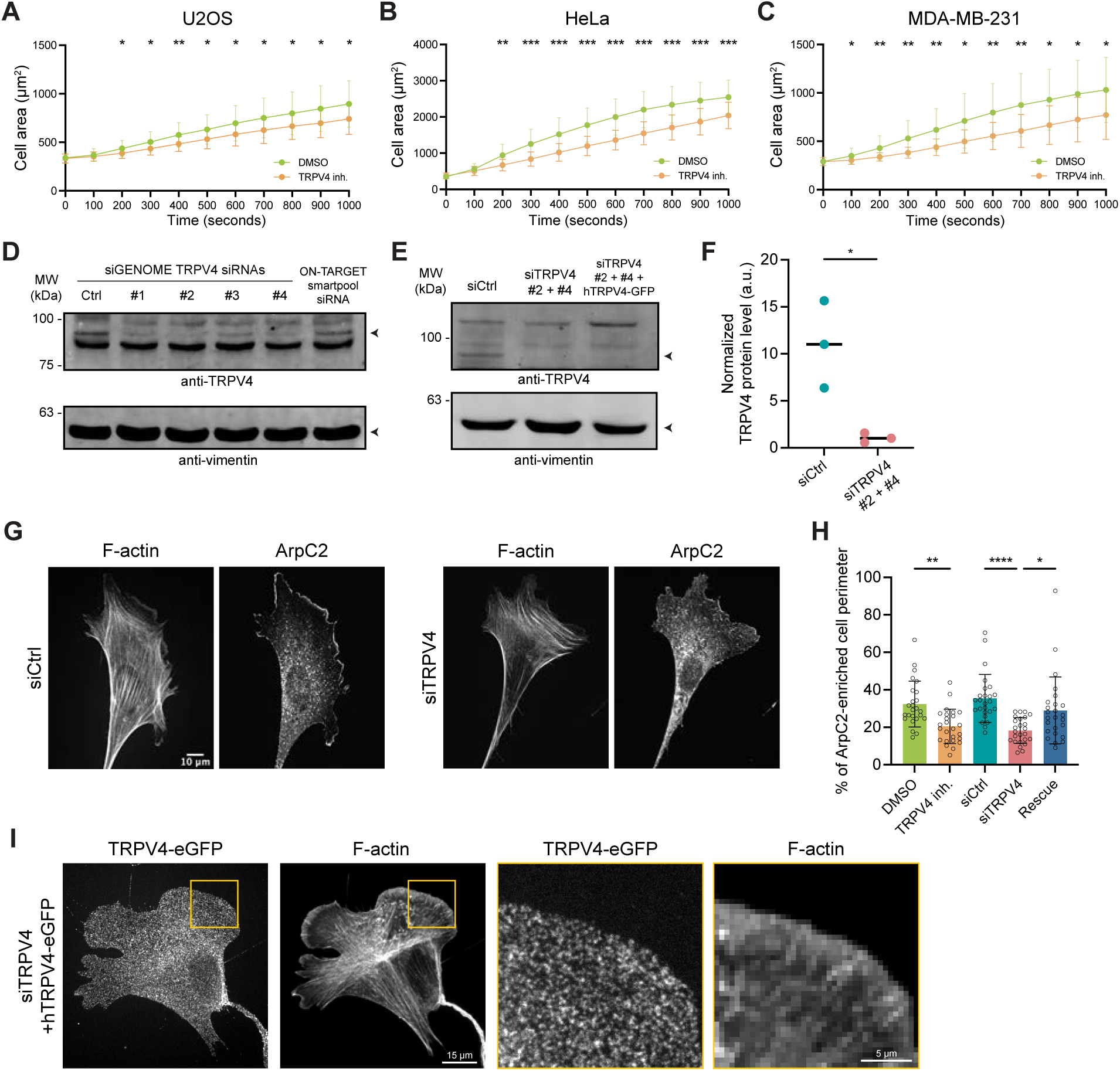
Suppression of TRPV4 impedes cell spreading across different cell types and disrupts lamellipodia formation. Related to Figure 1. (A-C) Quantifications of (A) U2OS, (B) HeLa and (C) MDA-MB-231 cell spread area over time in the presence of DMSO or TRPV4 inhibitor. n = 20 cells per condition. (D-F) Western blot analysis of endogenous TRPV4 level in siCtrl, siTRPV4, and rescue treatments 24 hr post-transfection. (D) Western blots showing the TRPV4 knockdown efficiencies of individual siRNAs and (E) a combination of TRPV4 siRNAs #2 and #4. (F) Quantifications of TRPV4 knockdown efficiency. n = 3 biological repeats. (G-H) Analysis of percent of ArpC2-enriched cell edge upon TRPV4 suppression. (G) Fluorescent phalloidin and anti-ArpC2 images of cells treated with 0.1% DMSO and TRPV4 inhibitor. (H) Quantifications of the percent of ArpC2-enriched cell edge. n > 20 cells per condition. (I) Immunofluorescence images of a TRPV4 siRNA treated MEF expressing hTRPV4-eGFP. The cell was stained for phalloidin (F-actin) and anti-GFP (hTRPV4-eGFP). Mann-Whitney test is used for comparisons between 2 treatments and one-way ANOVA followed by Tukey-Kramer post-hoc test for comparisons between 3 or more treatments. The data are presented as mean ± SD unless otherwise specified. * p < 0.05, ** p < 0.01, *** p < 0.005, **** p < 0.001.

**Figure S3.**
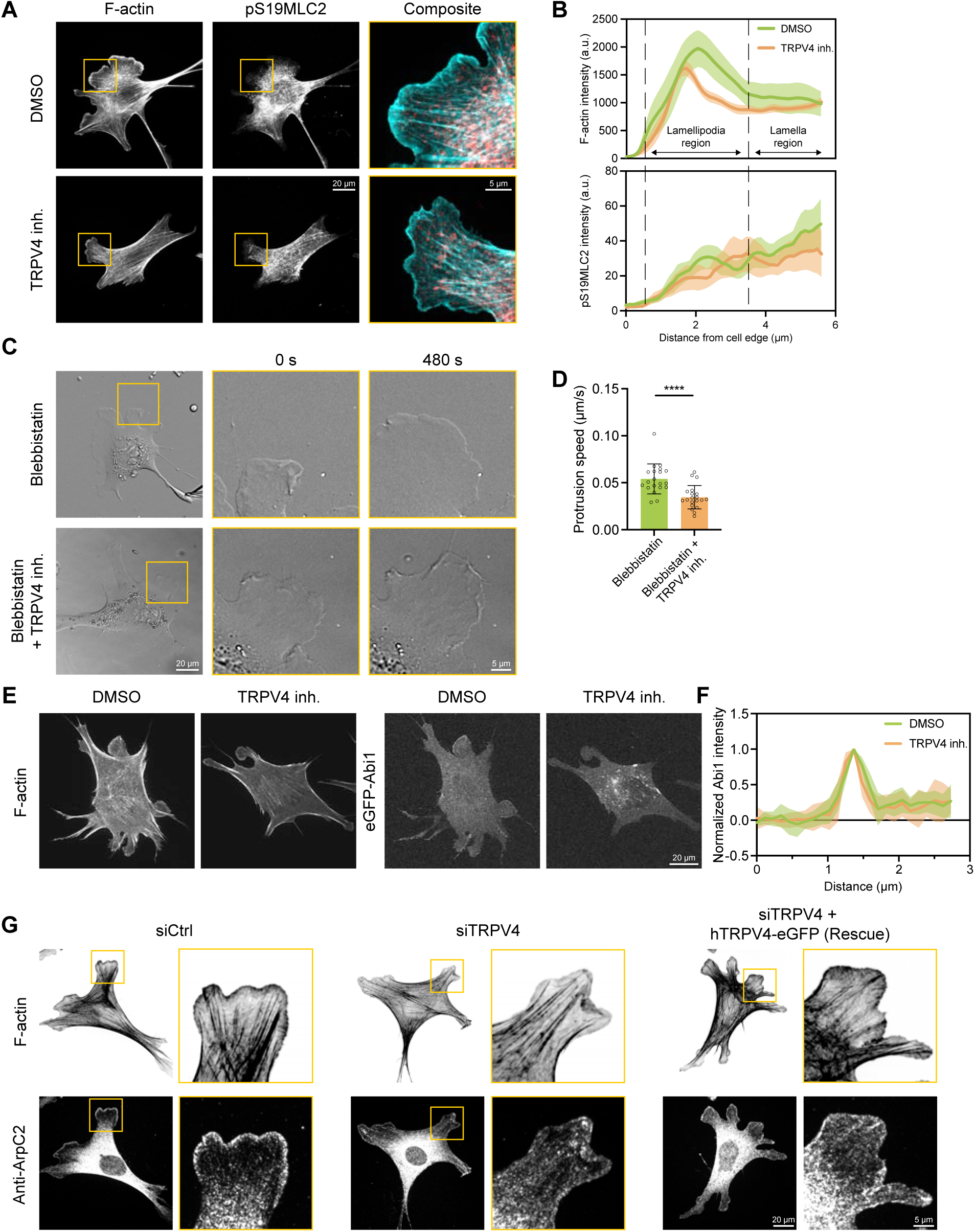
Myosin-II and the upstream regulators of Arp2/3 are not involved in the control of lamellipodia protrusive activity by TRPV4. Related to Figure 3. (A-B) Analysis of the effects of TRPV4 inhibition on phospho-myosin light chain (pS19MLC2). (A) F-actin and phospho-myosin light chain (pS19MLC2) images of MEFs treated with 0.1% DMSO and TRPV4 inhibitor (10 μM HC-067047). (B) Average line intensity profiles of the (top) F-actin and (bottom) pS19MLC2 staining near the cell edge in the presence of DMSO or TRPV4 inhibitor. n ≥ 6 cells per condition. (C-D) Analysis of the effects of myosin-II inhibition on lamellipodia protrusion speed. (C) DIC images of (left) blebbistatin-treated and (right) TRPV4 inhibitor-and-blebbistatin-co-treated MEFs. The yellow box indicates the region of the zoom-in shown on the right. (D) Quantifications of their protrusion speed. n = 20 cells per condition. (E-F) Localization of Abi1 in TRPV4 inhibited MEFs. (E) Fluorescent images of (left) F F-tractin-emiRFP670 and (right) eGFP-Abi1. (F) Average line intensity profiles of eGFP-Abi1 near the cell edge in the presence of DMSO or TRPV4 inhibitor. n = 9 cells per condition. (G) Fluorescent phalloidin and anti-ArpC2 images of MEFs treated with non-targeting siRNA (siCtrl), TRPV4 siRNA and TRPV4 siRNA + human TRPV4-eGFP (siRNA-resistant). Mann-Whitney test is used for comparisons between 2 treatments and one-way ANOVA followed by Tukey-Kramer post-hoc test for comparisons between 3 or more treatments. The data are presented as mean ± SD unless otherwise specified. * p < 0.05, ** p < 0.01, *** p < 0.005, **** p < 0.001.

**Figure S4.**
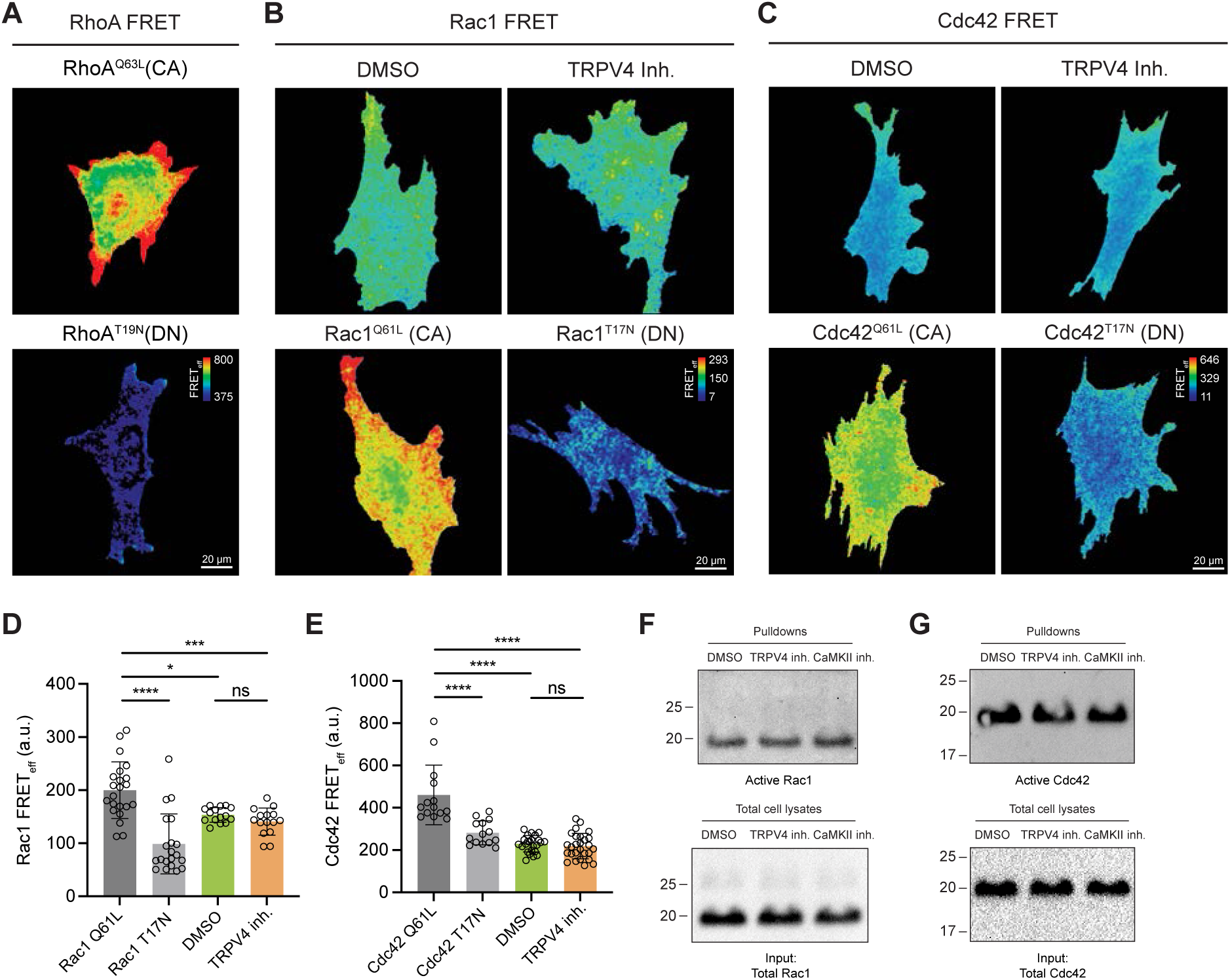
TRPV4 does not regulate the activity of Rac1 or Cdc42. Related to Figure 4. (A) Fluorescence images of RhoA FRET biosensor activation in MEFs expressing constitutively active (CA) or dominant negative (DN) RhoA mutant biosensors (B-E) Analyses of the effects of TRPV4 inhibition on Rac1 and Cdc42 activity by Förster Resonance Energy Transfer (FRET) imaging. MEFs were transiently expressing a Rac1 or Cdc42 FRET biosensor. Fluorescence images of (B) Rac1 and (C) Cdc42 FRET biosensors activation in cells treated with 0.1% DMSO or 10 μM HC-067047. Quantifications of the global (D) Rac1 and (E) Cdc42 FRETeff. n ≥ 14 cells per condition. (F-G) Western blot analysis of the activity of (F) Rac1 and (G) Cdc42 in MEFs treated with 0.1% DMSO, TRPV4 inhibitor (10 μM HC-067047), or CaMKII inhibitor (10 μM KN-93) by using the active GTPase pull-down assay. Mann-Whitney test is used for comparisons between 2 treatments and one-way ANOVA followed by Tukey-Kramer post-hoc test for comparisons between 3 or more treatments. The data are presented as mean ± SD unless otherwise specified. * p < 0.05, ** p < 0.01, *** p < 0.005, **** p < 0.001.

**Figure S5.**
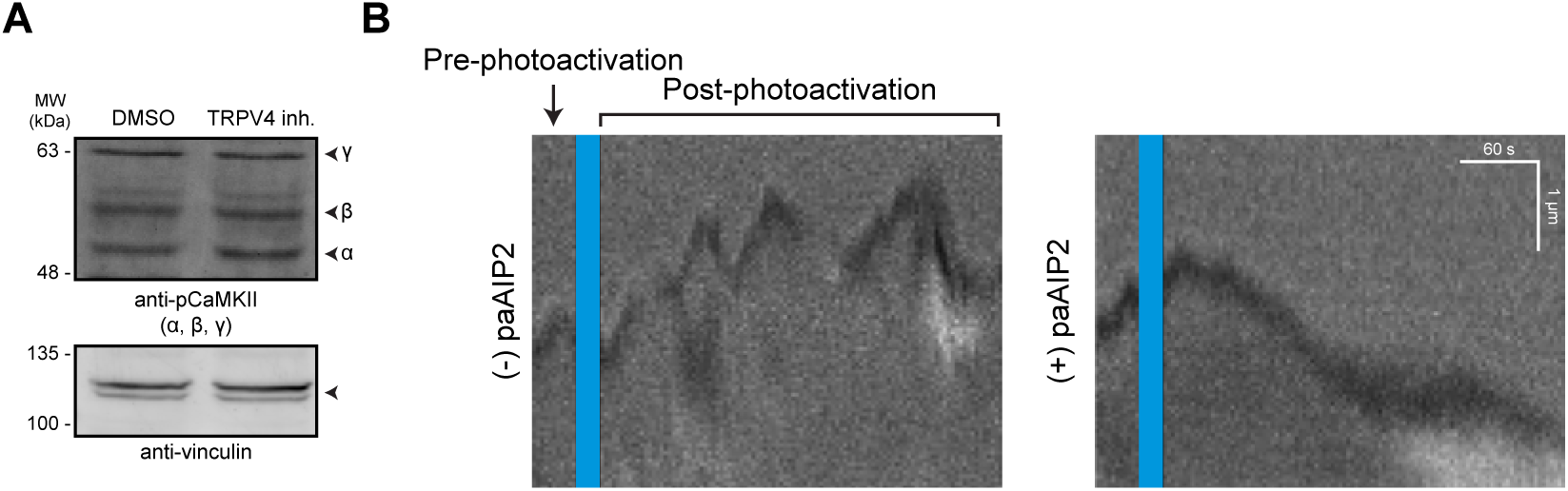
Local suppression of CaMKII perturbs lamellipodia protrusion dynamics. Related to Figure 5. (A) Western blot analysis of the level of phospho-CaMKII in MEFs treated with 0.1% DMSO or TRPV4 inhibitor (10 μM HC-067047). 3 isoforms of CaMKII (α, β and γ) are expressed. Vinculin was used as a loading control. (B) DIC Kymographs generated from the cells in the optogenetic CaMKII inhibition experiment. The kymographs show cell edge dynamics before and after photo-activation. The blue stripe indicates the photo-illumination period. They were extracted from cells without paAIP2 or with paAIP2 expression.

**Figure S6.**
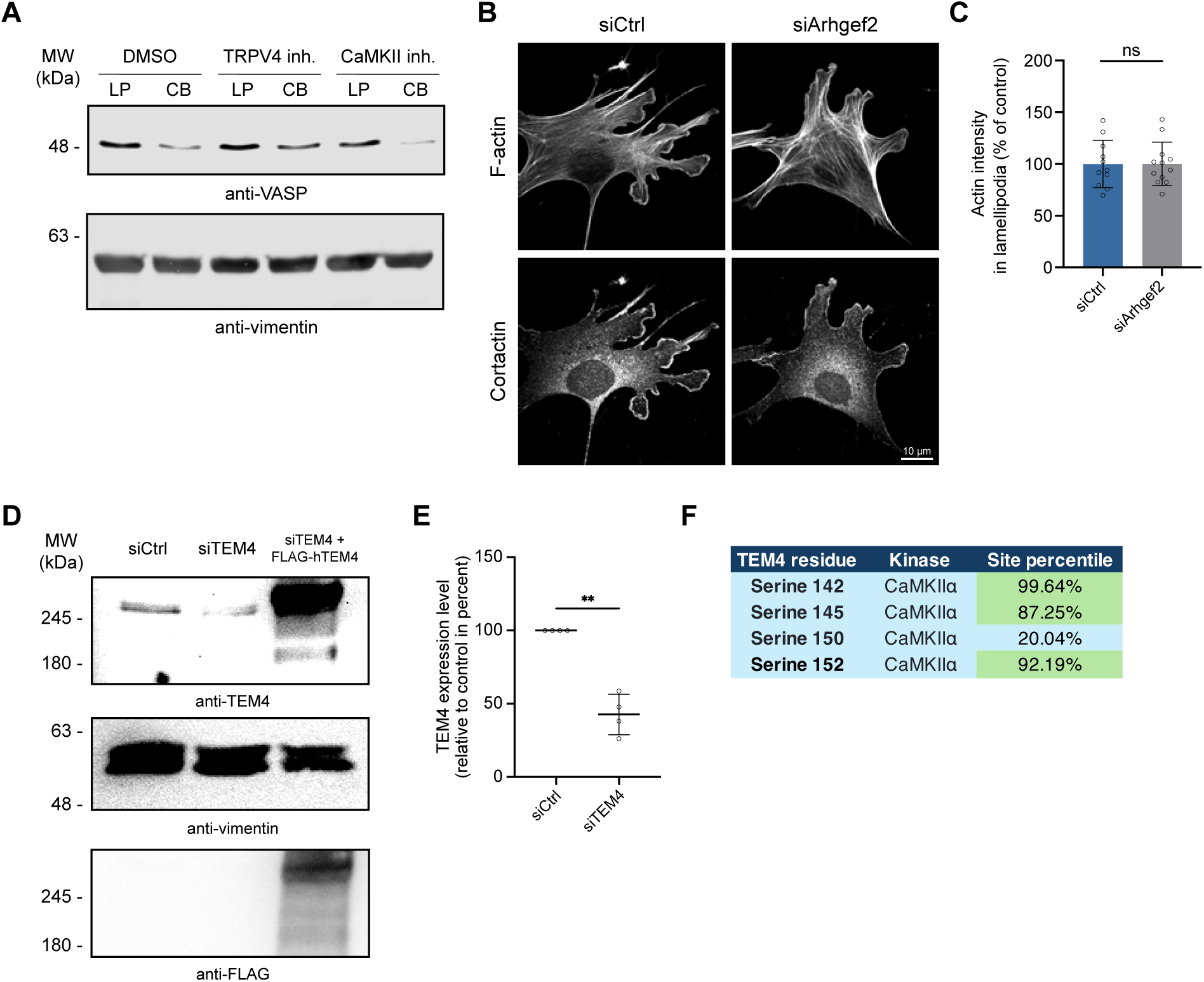
Arhgef2/GEF-H1 does not regulate lamellipodial F-actin. Related to Figure 6. (A) Western blot analysis of VASP (a lamellipodia marker) in purified lamellipodia (LP) and cell body (CB) fractions from MEFs migrating across a 3.0-μm pore size polycarbonate membrane. Prior to the fractionation, MEFs were treated with 0.1% DMSO, TRPV4 inhibitor (10 μM HC-067047), or CaMKII inhibitor (10 μM KN-93). Vimentin was used as a loading control. (B-C) Analysis of the effects of Arhgef2/GEF-H1 depletion on F-actin. (B) Fluorescent phalloidin and anti-cortactin images of MEFs treated with non-targeting siRNA (siCtrl) and Arhgef2/GEF-H1 siRNA (siArhgef2). (C) Quantifications of lamellipodial F-actin intensity upon Arhgef2 depletion. n = 10 cells per condition. (D-E) Western blot analysis of endogenous TEM4 level in siCtrl, siTEM4, and rescue treatments 72 hr post-transfection. (D) Western blots showing the TEM4 knockdown efficiency of the smart pool TEM4 siRNA and the over-expression of 3×FLAG-hTEM4 in TEM4-depleted MEFs. Vimentin was used as a loading control. (E) Quantification of TEM4 knockdown efficiency. n = 4 biological repeats. (F) Kinase prediction analysis of TEM4. The table shows the phosphorylation site percentile of TEM4 at serine residues 142, 145, 150 and 152 by CaMKIIα. Score percentiles that are above 80% were highlighted in green. The analysis was performed by using the PhosphoSite Plus database (www.phosphosite.org/uniprotAccAction?id=Q96PE2)^98^ . Mann-Whitney test is used for comparisons between 2 treatments. The data are presented as mean ± SD unless otherwise specified. * p < 0.05, ** p < 0.01, *** p < 0.005, **** p < 0.001.

**Figure S7.**
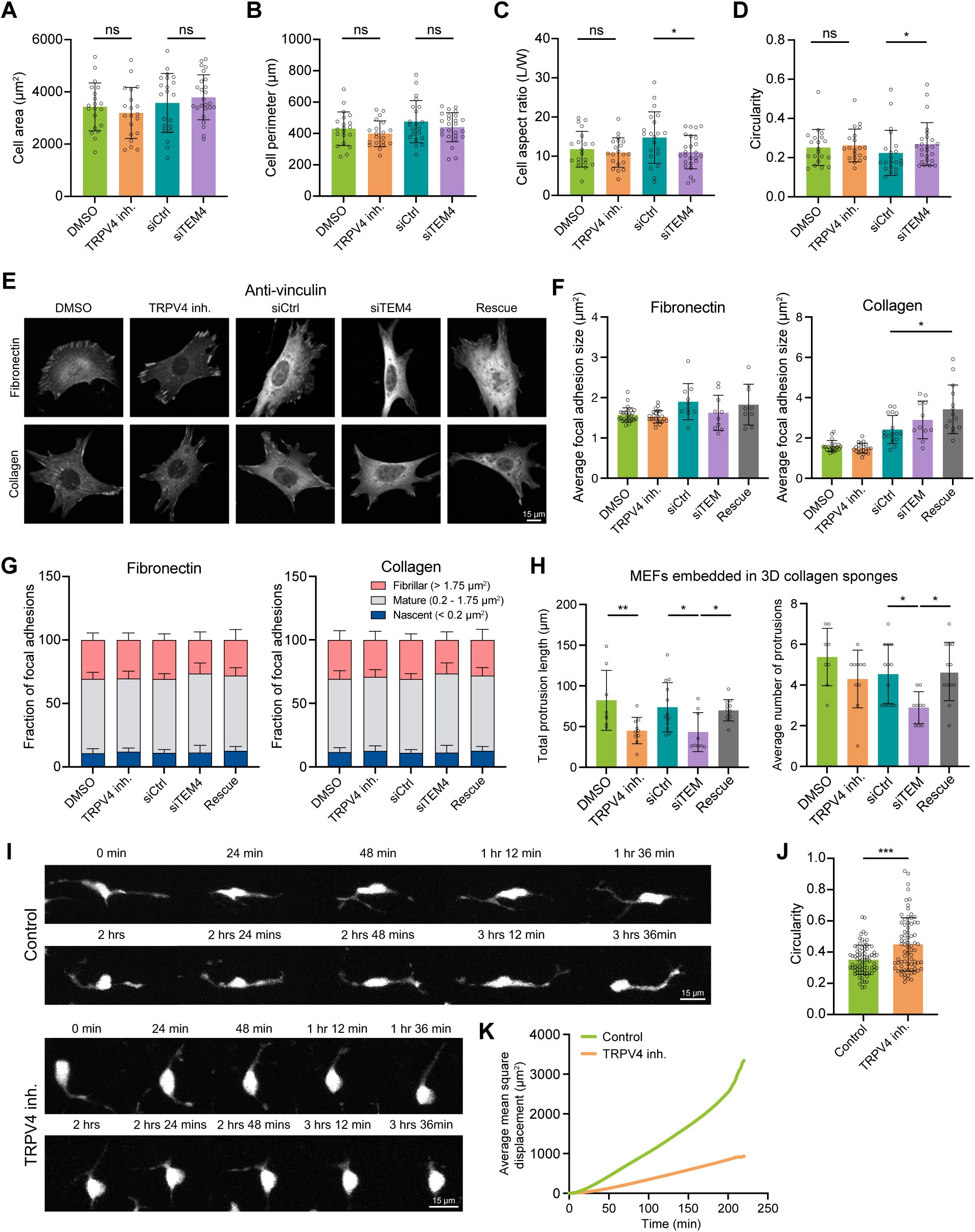
Suppression of TRPV4 signaling axis reduces cell protrusive activity while having a minimal effect on cell-ECM interaction and cell morphology. Related to Figure 7. (A-D) Cell morphological analysis of MEFs plated on a FN-coated coverslips upon TRPV4 inhibition or TEM4 depletion. n ≥ 20 cells per condition. (E-G) Analysis of the effects of TRPV4 inhibition and TEM4 depletion on focal adhesions. (E) Fluorescence images of MEFs stained for vinculin. Cells were treated with 0.1% DMSO, TRPV4 inhibitor (10 μM HC-067047), depleted for TEM4 or TEM4-rescued. Cells were plated on (top) fibronectin- or (bottom) collagen-coated glass coverslips. (F) Quantifications of the average focal adhesion size per cell on (left) fibronectin- and (right) collagen-coated glass coverslips. (G) Distribution of focal adhesion size (left) on fibronectin- and (right) collagen-coated substrates. n ≥ 10 cells per condition. (H) Quantifications of (left) the total protrusion length and (right) the average number of protrusions of MEFs embedded in a 3D collagen matrix. n ≥ 10 cells per condition. (I-K) Morphological and migration analysis of melanoblasts. Melanoblasts were imaged in *Mitf-Cre::Rosa26-floxed stop-tdTomato* E15.5 mouse embryo. (I) Timelapse images of melanoblast cell expressing tdTomato in (top) control and (bottom) TRPV4 inhibition. (J) Quantification of melanoblast cell circularity in the presence of TRPV4 inhibitor. (K) Quantification of the average mean square displacement (MSD) upon TRPV4 inhibition. n ≥ 74 cells per condition. Mann-Whitney test is used for comparisons between 2 treatments and one-way ANOVA followed by Tukey-Kramer post-hoc test for comparisons between 3 or more treatments. The data are presented as mean ± SD unless otherwise specified. * p < 0.05, ** p < 0.01, *** p < 0.005, **** p < 0.001.

## SUPPLEMENTAL VIDEO LEGENDS

### Video S1. Lamellipodial protrusion upon TRPV4 suppression. Related to Figure 1

MEFs transfected with siCtrl, siTRPV4 or siTRPV4 and eGFP-hTRPV4 (siRNA resistant TRPV4) migrating on 2.5 μg/mL FN-coated coverslips. DIC images were captured every 3 sec for 3 min 45 sec at 37°C. Top panels: Zoom-in views of a lamellipodial protrusion; Bottom panel: The entire migrating MEFs. The yellow box indicates where the zoom-in views was selected.

### Video S2. Calcium changes in lamellipodia before and after TRPV4 inhibition. Related to Figure 2

MEFs transfected with mCherry-T2A-GCaMP6f and F-tractin-emiRFP670 migrating on 2.5 μg/mL FN-coated coverslips. Multi-color fluorescence images were captured every 10 sec for 5 min 40 sec at 37°C. The cells were cultured in 0.1% DMSO supplemented phenol-red free DMEM in the first 1 min 50 sec and in 10 μM TRPV4 inh. supplemented phenol-red free DMEM after 2 min. Left panel: Timelapse images of ratiometric GCaMP6f/mCherry. Warmer colors indicate a higher Ca^2+^ level; Right panel: Timelapse images of F-tractin-emiRFP670 of a lamellipodial protrusion.

### Video S3. NP-EGTA-AM-induced lamellipodial protrusion. Related to Figure 2

MEFs transfected with mCherry-T2A-GCaMP6f and F-tractin-emiRFP670 migrating on 2.5 μg/mL FN-coated coverslips. Multi-color fluorescence images were captured every 15 sec for 30 sec before Ca^2+^ uncaging and for 2 min after the uncaging. The white box indicates the region of 405 nm photo-illumination. Top panels: F-tractin-emiRFP670 images of MEFs; Bottom panels: GCaMP6f images of MEFs. Brighter colors in the lookup table indicate a higher Ca^2+^ level; Left and Right panels: MEFs without and with NP-EGTA-AM.

### Video S4. Optogenetic inhibition of CaMKII in lamellipodia. Related to Figure 5

MEFs transfected with histone H2B-eBFP2 (Left; (-) paAIP2) or histone H2B-eBFP2-P2A-paAIP2 (Right; (+) paAIP2) were seeded in 2.5 μg/mL FN-coated glass-bottom dishes. Timelapse DIC images were acquired every 3 sec for 36 sec before and 5 min after 405 nm photo-illumination (3 sec). The purple box indicates the region of photo-illumination.

### Video S5. Effects of TRPV4 inhibition and TEM4 depletion on 2D migration. Related to Figure 7

MEFs transfected with histone H2B-mEmerald migrating on 2.5 μg/mL FN-coated glass coverslips. Top panels: MEFs treated with 0.1% DMSO or 10 μM TRPV4 inh. Bottom panels: MEFs co-transfected with siCtrl or siTEM4. For each condition, timelapse DIC and fluorescent histone H2B images were acquired every 12 min. The color code on the histone H2B timelapse images represents the time.

### Video S6. Effects of TRPV4 inhibition on melanoblast migration. Related to Figure 7

Melanoblast migration in E15.5 trunk skin explants (left) without or (right) with 10 μM TRPV4 inh. To label the embryonic melanoblasts, a melanocyte-specific *Mitf-Cre* transgene was used to induce tdTomato expression from a *Rosa26-floxed stop-tdTomato* fluorescent reporter. Confocal z-stacks were taken every 2 min continuously for 4 hr.

